# Climate structures global associations between human footprint and arthropod diversity

**DOI:** 10.64898/2026.05.29.728852

**Authors:** Alessandro Zito, Tommaso Rigon, Tomas Roslin, Pekka Niittynen, Paul D. N. Hebert, Evgeny V. Zakharov, Sujeevan Ratnasingham, iBOL Consortium, Otso Ovaskainen, David B. Dunson

**Affiliations:** Department of Biostatistics, Harvard University, Boston, MA, U.S.A; Department of Economics, Management and Statistics, University of Milano-Bicocca, Italy; Department of Agricultural Sciences, University of Helsinki, Helsinki, Finland; Water, Energy and Environmental Engineering Research Unit, University of Oulu, Oulu, Finland; Centre for Biodiversity Genomics, University of Guelph, Guelph, Canada; 150 Research Lane, Guelph, Canada; Department of Biological and Environmental Science, University of Jyväskylä, Jyväskylä, Finland; Department of Statistical Science, Duke University, Durham, NC, U.S.A

## Abstract

Human pressure is widely regarded as a major driver of insect decline, yet whether its impact is globally uniform remains unknown. This is because most arthropod species are undescribed, available diversity metrics often disagree and are sensitive to sampling efforts, and many existing analyses combine heterogeneous sampling designs. Using data on 11.5 million arthropods collected across all continents via standardized protocols, we find that the association between human footprint and arthropod diversity is structured by climate: richness is 17%, 19%, and 24% lower at heavily impacted sites in temperate, tropical, and dry regions, and increases in polar ones. These patterns are revealed by Hubbell regression, a novel statistical model that enables reliable comparisons across unequal sampling efforts in terms of familiar biodiversity indicators. Together, the data and the method make the human imprint on undescribed biodiversity measurable at a global scale.

---

Global biodiversity is in rapid decline [1], with current extinction rates surpassing anything seen for the past 66 million years [2]. Such massive loss threatens essential ecosystem functions [3] and services [4], with substantial economic and health repercussions [5–7]. To counter this crisis, we urgently need a sound understanding of the patterns and drivers underpinning global biodiversity. However, three features make biodiversity science especially challenging.

First, most biodiversity remains uncharacterized. Of the estimated 8.7 million eukaryotic species on Earth, only about 1.2 million have been described by science [8]. Second, community ecologists lack consensus on which biodiversity metric to use to measure change. For example, the number of proposed metrics to date exceeds 500, leading to conflicting conservation priorities even for the same sites [9]. To be effective, a useful biodiversity metric should be scientifically grounded yet policy-relevant, sensitive to change across scales and taxa, and informative about the state of nature alongside the pressures shaping it [10]. Third, a general analytical framework for pinpointing the drivers of variation in local and regional biodiversity remains lacking [11, 12].

These challenges come to a head amid ongoing losses in arthropod populations [13, 14]. While long-term studies suggest that terrestrial arthropod species richness is in sharp decline and that these disruptions are linked to anthropogenic impacts [15], available data are dominated by a minority of well-known insect taxa, with strong taxonomic and geographic biases and limited consideration of undescribed taxa [14]. As a result, global studies have relied on heterogeneous assemblages compiled across separate sources [16–18], rather than on standardized sampling protocols. To accurately resolve the scale and variation of the human imprint on arthropod diversity, we need a step change in both the *scale* of the data and the *approach* to measuring and comparing biodiversity.

In this paper, we address both needs by providing a globally curated arthropod dataset and a statistical method to analyze it. Specifically, we present systematic samples of 11.5 million arthropod individuals collected by the Global Malaise Trap Program (GMTP; [19]) in 15,176 sampling events across 1,186 sites spanning all continents (Figure 1a,b). These data were collected using a strictly standardized protocol over a near-twenty-year collection period, and represent the largest global effort to characterize species-rich communities one individual at a time. Then, we introduce a single coherent biodiversity measurement framework to enable comparisons across samples of different sizes and coverage, and to examine their interplay with environmental conditions. Our proposed model, called *Hubbell regression*, is a novel generalized linear model for species richness that infers relationships between biotic and abiotic factors and the fundamental biodiversity number *α* [20, 21]. Unlike conventional richness regressions, which usually model observed counts or an extrapolated richness estimate, Hubbell regression models the species-discovery process conditional on the number of sampled individuals, preserving valid accumulation curves. In particular, *α* controls how quickly new (including undescribed) species are discovered as additional individuals are sampled while also accounting for unequal sampling efforts in a statistically principled manner. Moreover, several diversity metrics, such as the Shannon, Simpson, and Tsallis indices, can be recovered from *α* within the Bayesian species sampling model framework [22, 23]. The complete workflow is described in Figure 1c.

**Figure 1.**
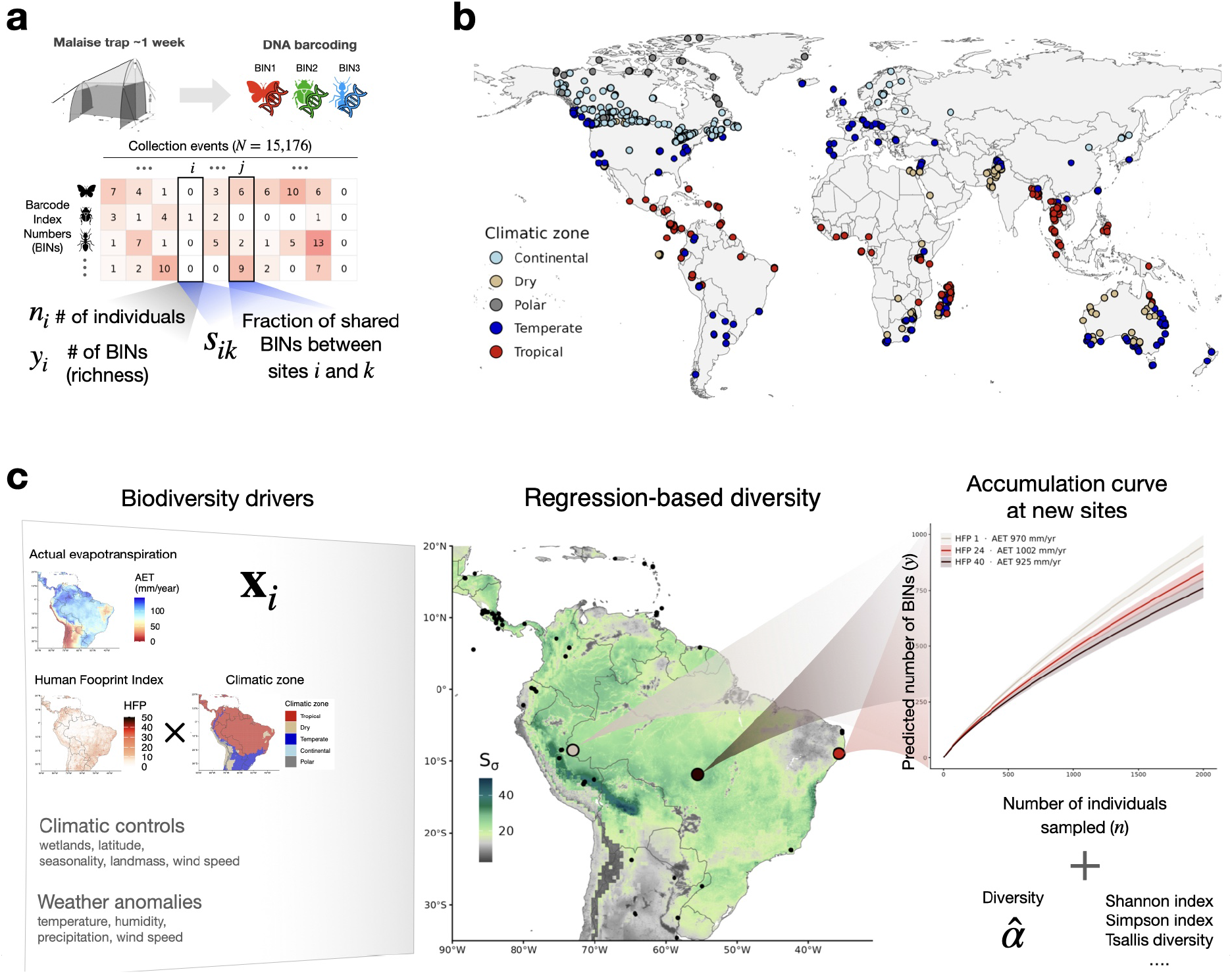
GMTP study design and details of the Hubbell regression. (**a**) Description of the data. In each site, specimens are collected for approximately one week and later classified into BINs using DNA barcoding methods. For each sample, we record the number of specimens *n*_*i*_, the sample richness *y*_*i*_ (i.e., number of unique BINs), and the fraction of shared BINs between every two locations. (**b**) Locations of the sampling sites and their climatic classification according to the Köppen–Geiger zone. (**c**) End-to-end illustration of the analysis with the Hubbell regression. We denote covariates for location *i* as **x**_*i*_. We use AET and the interaction between HFP and climatic zone as primary drivers for arthropod diversity, and further control for climate and weather. This allows us to predict regression-based diversity *S*_*σ*_ (restricted here to the Amazon rainforest for illustration). Black points denote GMTP collection sites, while colored points correspond to hypothetical new sites, chosen to have similar values of evapotranspiration, latitude, wind speed, wetland cover, and biogeographic realm, and to differ mainly in human footprint. Hubbell regression can extract accumulation curves at any new location based on environmental characteristics, and further predict many more biodiversity indices.

Using Hubbell regression and the GMTP data, we find that the association between human pressure and arthropod richness is structured by climate [24–27], shifting from a 17–24% decline in temperate, tropical, and dry regions to an increase in polar areas. Hence, the human imprint emerges not as a uniformly negative force but as a climate-structured phenomenon. Our results are available at the HubbellGLM Atlas (https://hubbellglm.org/), an interactive platform that enables scenario evaluations and predictions to inform future studies and conservation efforts.

## Hubbell regression: a model to relate biodiversity to environmental features

The characterization of biodiversity using diversity indices has a long history in statistics, with early contributions dating back nearly a century [20, 28–33]. Most such metrics are primarily intended to provide specific perspectives on the diversity of individual sites. However, how to incorporate environmental features into familiar ecological tools, such as accumulation curves and species richness, remains unclear. The problem is compounded by the fact that many metrics are sensitive to sampling effort and evenness. We now propose a model that overcomes these limitations.

Our starting point is the statistical machinery behind the popular unified neutral theory of biodiversity and biogeography [21, 34]. At its core, the unified neutral theory postulates a *fundamental biodiversity number*, which we refer to as *α*, that controls how quickly new species are discovered as additional individuals are sampled. In particular, *α* governs the species richness, i.e., the number of unique species *Y* ^(*n*)^ observed among *n* = 1, 2, … individuals, via the distribution

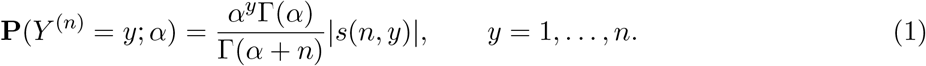

The quantities |*s*(*n, y*)| are called signless Stirling numbers of the first kind, and Γ(*x*) is the gamma function. Equation 1 follows directly from the Ewens sampling formula [35, 36], which describes species abundances under the neutral theory in a sample undergoing pure ecological drift. Importantly, many familiar biodiversity indices, such as Shannon and Simpson diversity, can be expressed as direct functions of Hubbell’s *α* [22, 23, 37]; we report formulas in Table S1. Therefore, knowing *α* means also knowing a broad range of additional biodiversity metrics.

To further estimate the relationship between environmental factors and *α* across multiple collection sites, we introduce a novel statistical framework, which we call *Hubbell regression*. Our approach is as follows. First, we model species richness according to Equation 1. This implies that the mean of the number of species, denoted as *µ*, is equal to

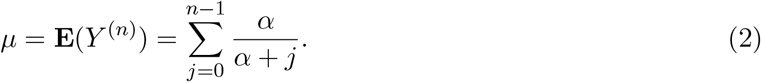

Second, we link the mean *µ* and a set of *p* environmental features **x** ∈ ℝ^*p*^ via a function *g* as

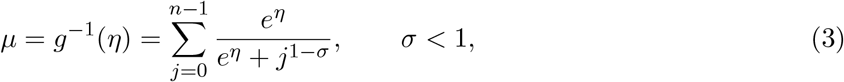

where *η* = **x**^⊤^***β*** is a linear predictor, ***β*** are regression coefficients capturing the relationship between the covariates and *µ*, and *σ* is a parameter controlling the growth rate of the species richness with *n*. Third, to account for potential dependence among observations arising from common species across sites, we borrow from ideas in spatial econometrics [38] and correct standard errors using the Jaccard similarity index between each pair of samples. This yields robust *P* values when assessing the significance of the impact of any biodiversity driver (Figure S1). In Materials and Methods, we show that Hubbell regression belongs to the statistical class of generalized linear models [39], and therefore enjoys several important properties and simple estimation algorithms; see Algorithm S1.

Both Equation 2 and Equation 3 ensure that *µ* stays between 1 and *n*, leading to well-behaved in-domain predictions. In particular, Equation 2 grows logarithmically in *n*, while Equation 3 encompasses a broader range of growth rates depending on *σ*. If *σ* = 0, we retrieve Hubbell’s *α* as *α* = *e*^*η*^ = exp*{***x**^⊤^***β****}*. Values of *σ* ∈ (0, 1) yield a polynomially growing curve, while values *σ <* 0 yield a curve converging to a finite value as *n* goes to infinity (Figure S2). We summarize these behaviors via the single quantity *S*_*σ*_(*η*), called the *regression-based diversity index*, which we define as the limit of the ratio between the link function *g*^−1^(*η*) and the growth of the curve with *n* (Figure 1c). For example, when *σ* ∈ (0, 1), we have *S*_*σ*_(*η*) = *e*^*η*^*/σ* (Theorem S1). This lets us translate unit changes in *η* into percentage changes in diversity, as in a standard log-linear model.

## Climatic associations between human pressure and arthropod diversity

The GMTP is an ongoing collection effort to map global arthropod diversity. Figure 1a and b describe the data assembled here. We restrict the analysis to arthropods collected using Malaise traps between 2007 and 2024. We assigned individuals to species using DNA sequencing. Because most species in the samples lacked scientific names, we clustered sequences into Barcode Index Numbers (BINs) based on sequence similarity [40]. BINs are reliable proxies for species and units for biodiversity assessment [40, 41]. Therefore, we measure species richness as the number of unique BINs found at a site. In total, these samples yielded 691,283 unique BINs.

The broad coverage and standardization of the GMTP data allow investigation of arthropod diversity at the global scale. While human presence is widely regarded as one of the key drivers of recent insect decline [13, 16, 17], such pressure never acts in isolation: its imprint is superimposed on a climatic backdrop that sets the energetic baseline of every community [18]. To test this hypothesis, we estimate a sequence of nested models of increasing environmental complexity where the human footprint index (HFP; [42]) is interacted with the broad Köppen–Geiger climatic zone of the site (Continental, Dry, Polar, Temperate, and Tropical) [43], further controlling for the general imprint of environmental energy (measured via *actual evapotranspiration*; AET, [24]), and additional geographic, climatic, weather, and seasonal variables (models M0–M7, Materials and Methods). We account for unmeasured spatial and temporal dependence by correcting standard errors via the Jaccard similarity index, defined as the fraction of shared BINs between each pair of samples. This is inversely related to the geographic distance between sites (Figure S3).

We find that the imprint of human activity on arthropod biodiversity varies substantially and systematically across the globe. Using Hubbell regression, we estimate that an increase of 5 units on the HFP scale is associated with a 5.3% reduction in diversity in tropical areas (*β* = −0.011; *P <* 10^−18^), a 5.6% reduction in dry zones (*β* = −0.012; *P <* 10^−7^) and a 3.8% reduction in temperate zones (*β* = −0.008; *P <* 10^−6^), while polar areas show an increase of 18.4% (*β* = 0.034; *P <* 10^−8^). In continental areas, the association is not distinguishable from zero (*β* = 0.001; *P* = 0.55). Consistent with the hypothesis of [24], our analysis estimates that AET provides the largest incremental model-fit contribution among alternative environmental variables, including vapor pressure deficit, relative humidity, precipitation, and mean annual temperature (Table S2 and Figure S4). We estimate that a 100 mm/year increase in AET is associated with a 9.5% increase in diversity on average (*β* = 0.009; *P <* 10^−35^). These estimates are robust to including more controls in the specification (Table S3). Moreover, across random and spatially structured cross-validation schemes, Hubbell regression matched or improved predictive accuracy relative to conventional richness regressions while preserving admissible accumulation curves (Figures S5 and S6). Figure 2 illustrates the interplay between AET, HFP, and climatic zone for both the regression-based diversity index *S*_*σ*_(*η*) and the predicted species richness *ŷ*_*n*_, assuming a sample of *n* = 2000 individuals. Related extensions to other biodiversity indices are shown in Figure S7.

**Figure 2.**
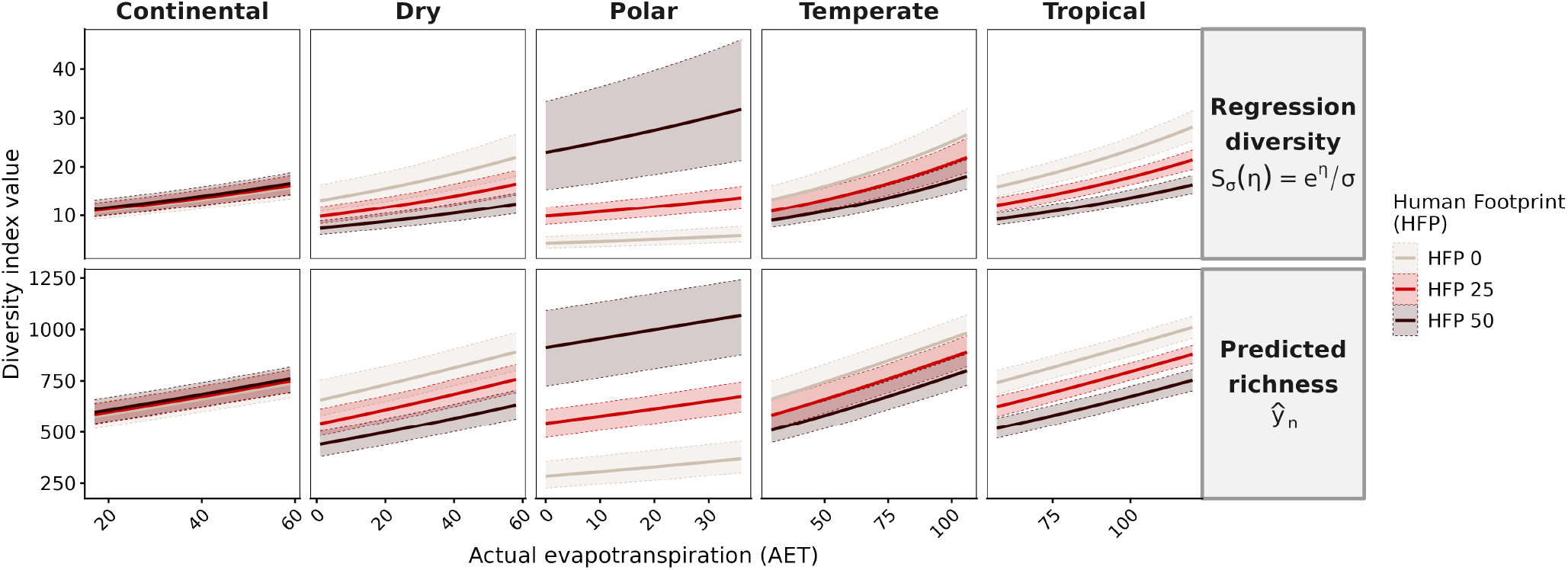
Variations of AET and HFP across climatic zones. All curves are predictions from the most complete specification for the year 2024, fixing the community size to *n* = 2000, *σ* = 0.5812 (the maximum likelihood value), the collection week to the 26th week of the calendar year, and the collection duration to 7 days. Latitude is set to the mean of each zone on the global grid; wetlands, wind speed, the residual temperature term, and all weather anomalies are set to zero, and the landmass and isolation terms are set to their sample medians. The realm is the most common realm of each zone on the global grid (Nearctic for Polar, Palearctic for Continental, Dry, and Temperate, Neotropic for Tropical). Bands around the lines indicate 95% bootstrap confidence intervals, computed using Jaccard-adjusted standard errors. The AET range shown for each zone is its 5th–95th percentile on the 2024 global gri.d

Our estimates can be used to explore alternative scenarios for changes in AET and HFP. To do so, we predict biodiversity and richness using a 0.1° *×* 0.1° grid (Figure 3, left panels). Assuming that a one-week collection during the 26th Julian week of the year (adjusted by six months if below the equator) captures a total of *n* = 2000 specimens anywhere on Earth (i.e., the mid-summer abundance peak), our analysis indicates that a median of 171 species (18.9% of potential richness) is predicted to be lower in tropical sites characterized by high human footprint (HFP *>* 30). These effects are larger in dry sites (153 missing species, 24.4% of potential richness) and smaller in temperate sites (114 species, 16.6%); see Figure 3a. Overall, a 5-unit intensification of human presence everywhere leads to up to a 5.1% reduction in potential richness (Figure 3b), while a 10% increase in AET is associated with an average 2.5% increase (Figure 3c). Peaks above 10% are observed in the tropical

**Figure 3.**
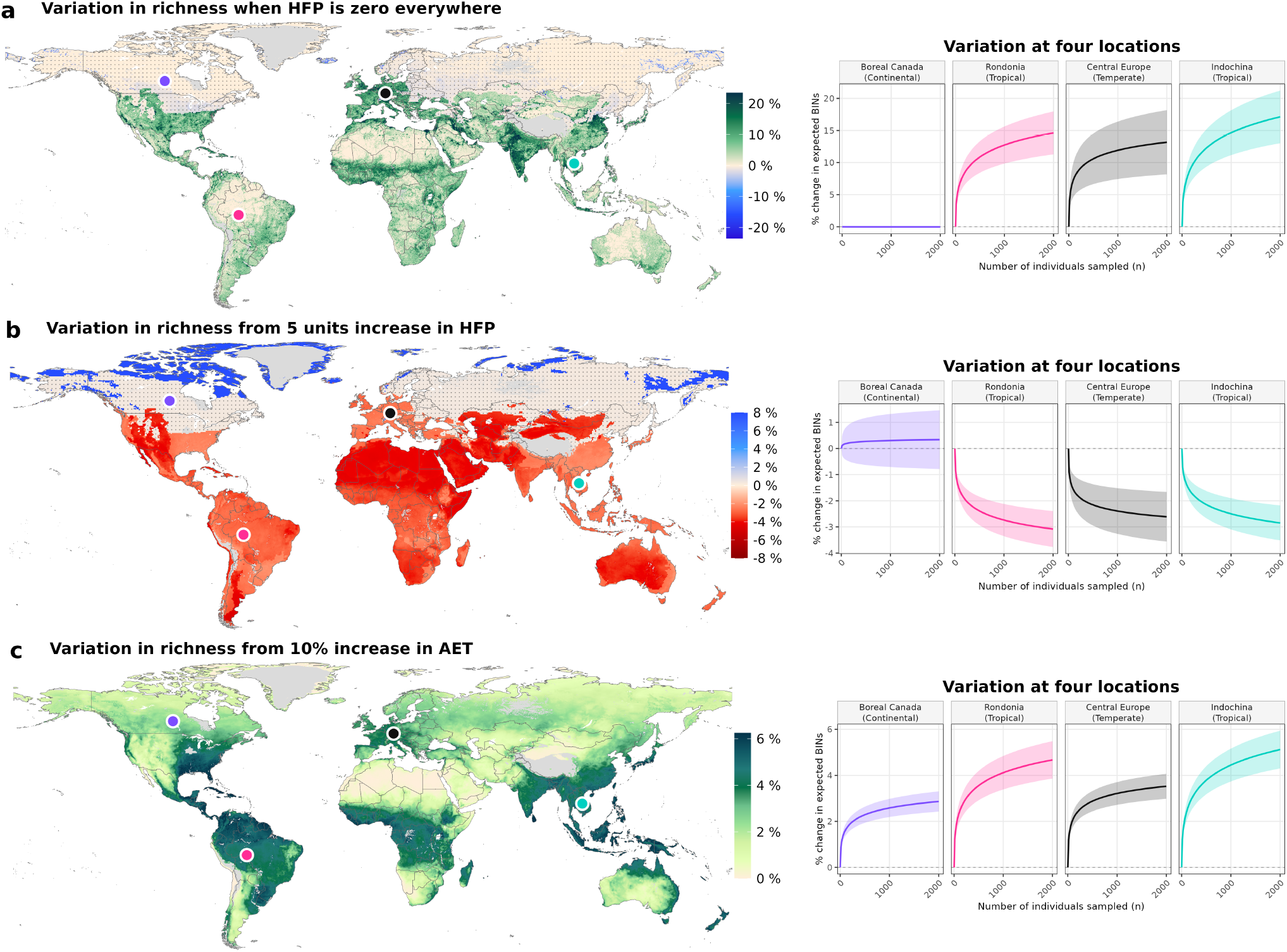
Global shifts in arthropod richness from variations in biodiversity drivers. Left plots denote the percentage variation in predicted average richness *Y* ^(*n*)^ assuming that *n* = 2000 specimens are captured at any location when HFP is set to zero (**a**), after a 5-unit increase in human pressure, capped at the maximum of 50 (**b**), and after a 10% increase in evapotranspiration (**c**). Values are calculated using the 26th week of the year, assuming 7 days of collection, with no weather anomalies, for covariate values in 2024. Gray values indicate areas where extrapolations are unreliable. Cells whose 95% interval covers zero (primarily continental areas) are stippled, applied to 2° blocks in which more than half the cells are non-significant. Predictions are obtained by letting *σ* = 0.5812, the maximum likelihood value. Right plots: percentage difference in accumulation curves at four locations. Confidence intervals use Jaccard-adjusted standard errors.

Caribbean and Central America (Figure 3c). The right-hand panels of Figure 3 show how the same scenarios lead to variations in accumulation curves at four out-of-sample locations. Removing human pressure leaves a boreal Canadian cell (HFP = 0.3) unchanged (−0.02% at *n* = 2000), while recovering 14.6% of richness in Rondônia, 13.2% in central Europe, and 17.1% in Indochina (HFP = 23.2, 24.2, and 29.8). These deficits depend on sampling effort: at *n* = 44 individuals, the three impacted cells are indistinguishable (5.3–5.9%), separating only as *n* grows. Human pressure therefore flattens the accumulation curve rather than removing a fixed proportion of species, so the magnitude of the estimated loss is itself a function of community size.

## Discussion

In this paper, we have leveraged a set of standardized arthropod samples collected globally to show that arthropod biodiversity is systematically associated with the human footprint, and that this association varies with climate and productivity. In tropical and temperate regions and in arid zones, a higher human footprint is associated with reduced insect diversity, while in polar areas, an increase in the human footprint is associated with increased diversity. These contrasting effects may reflect narrow thermal and ecological niches presumed characteristic of tropical arthropods. Because tropical environments experience minimal seasonal temperature variation, such species are assumed to be highly specialized and particularly vulnerable to microclimatic disruptions [44–46]. Conversely, the positive association between HFP and arthropod diversity in polar regions may highlight the role of human infrastructure as a source of artificial heating and direct biomass subsidies, thereby facilitating species’ survival and enabling the range expansion of temperate arthropods into polar latitudes [47].

Importantly, three main advances contribute to the credibility of the current work: first, we draw on systematic sampling using standardized techniques. Second, we assign individuals to species using methods independent of geographic or taxon-specific variation in taxonomic expertise. Third, we rely on information inherent in the accumulation of new species within the data range, rather than on either observed or extrapolated species richness. The first perspective ensures appropriate comparisons and avoids both noise and bias typically associated with meta-analyses of data from mixed sources (e.g., [16]). From the second perspective, using DNA-based species delimitation criteria [40] allows us to comprehensively characterize global biodiversity across arthropod groups, circumventing the challenge that a large fraction of species remain undescribed to this day [48]. Lastly, using species discovery information within the data range improves the reliability of biodiversity estimates. This is because insect communities most often feature a (very) long tail of rare species [20, 49, 50]. Thus, realistically sized samples of local arthropod communities will typically fail to reveal true species richness, and asymptotic estimates of true richness will be subject to both bias and substantial uncertainty.

Where global environmental change calls for efficient metrics to characterize the current state of biodiversity and to attribute its change to relevant drivers [51], we have introduced a novel method that achieves both. Drawing on classic ecological theory [20, 21] and its connection to Bayesian statistics [37], we developed a principled regression framework to estimate current biodiversity and relate it to the local environment. This framework achieves predictive performance comparable to or better than that of existing methods (Figure S5), while offering a precise interpretation in terms of the fundamental biodiversity number *α*. This number can be converted into several other diversity indices, avoiding the need to choose among them. Hence, Hubbell regression meets the standards increasingly demanded of modern biodiversity metrics: it is theoretically principled, tractable, and sensitive across scales and taxa [9].

By comparing predicted species richness in a hypothetical community of 2000 individuals with the value predicted when the human footprint is set to zero, we estimate a deficit of 24.4% of potential richness (153 species) at heavily impacted dry sites (HFP *>* 30), 18.9% (171 species) at tropical sites, and 16.6% (114 species) at temperate ones, whereas continental sites show no deficit and polar sites exhibit species gain. These values exceed the global average 13.6% reduction in species richness attributed to land-use pressures by Newbold *et al*. [16], as expected when conditioning on the most heavily impacted locations rather than averaging over all land. Hence, our framework shows that the magnitude of these deficits depends on the underlying climatic baseline, with the most severe ones concentrated in dryland and tropical ecosystems. The same framework applies wherever standardized samples and environmental covariates are available, making such comparisons reproducible across taxa and regions.

## Acknowledgments

We are grateful to the researchers of the full Lifeplan research group for the many helpful discussions.

## Author contributions

A. Z. and T. Rig. developed the methodology and the associated theory, and performed all statistical analyses; A. Z., T. Rig., T. Ros., O. O., and D. D. wrote the paper; P. H., E. Z., S. R., and the iBOL Consortium designed and performed data collection for the GMTP. P. N. provided general guidance on statistical analyses and dataset assembly. S. R. and A. Z. developed the HubbellGLM Atlas at https://hubbellglm.org/.

## Data and code availability

Code to reproduce the figures and the analysis in the paper is available at the GitHub repository https://github.com/alessandrozito/HubbellGLM-paper/. All models in this paper are estimated via the R package HubbellGLM, available at the GitHub repository https://github.com/alessandrozito/HubbellGLM. The full extent of the data can be visualized at https://hubbellglm.org/.

## Supplementary Material

### Materials and methods

#### Mathematical foundations of biodiversity

The problem of characterizing biodiversity has deep roots in the history of statistics, with seminal contributions from Fisher [20] and Preston [29]. From the 1920s until today, a large body of work has aimed to estimate the number of species in the population from a finite sample [28, 30, 31, 33, 52], to characterize the relative dominance of different species in a community (evenness; [53, 54]), and to combine metrics of species richness and evenness into a single metric [32]. Ecologists have since adopted multiple metrics to describe global patterns in species richness [27] and to prioritize the most diverse sites for conservation.

The theoretical foundations of many biodiversity indices rest on Bayesian nonparametric species sampling models [22, 37]. The cornerstone of these models is the Dirichlet process [55, 56], which is mathematically equivalent to the Ewens sampling formula [35] that governs the distribution of taxon abundances in a community of size *n* in the unified neutral theory of biodiversity and biogeography [21, 36]. This equivalence allows us to express all biodiversity indices in terms of the fundamental biodiversity number *α*; see [23].

From an ecological perspective, a Dirichlet process assumes that the composition of a community is an infinite sum of species, written as 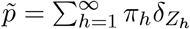. Here, *Z*_1_, *Z*_2_, … represent distinct species labels (e.g., BINs in the current study), sampled from a common baseline distribution *P*_0_. The parameters *π*_1_, *π*_2_, … are the relative abundances of each corresponding species label, with 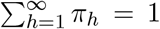. In particular, each probability *π*_*h*_ is constructed via a stick-breaking formulation: 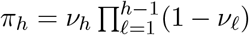, where *ν*_*h*_ ~ Beta(1, *α*) independently for each *h*, and *α* is again the fundamental biodiversity number.

The above construction allows us to express all biodiversity indices in a general way as 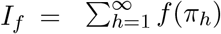, for some function *f* : (0, 1) → ℝ_+_ [57]. Moreover, the stick-breaking probabilities are in size-biased order, meaning that *π*_1_ is the relative abundance of the species of a randomly sampled individual, *π*_2_ that of the next new species to be discovered, and so on. As a consequence, the expected value of *I*_*f*_ simplifies elegantly to an expectation that depends only on the first sampled species:

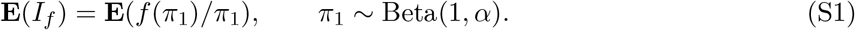

Letting *f* (*π*) = *π*^2^, *f* (*π*) = −*π* log *π*, and 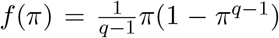 returns the model-based Simpson index, Shannon index, and (unnormalized) Tsallis number of order *q*, respectively. In these cases, Equation S1 is available analytically from properties of the beta distribution. While the relationship between the Simpson index and *α* has been established in the ecological literature [58], connections with the other indices have not been explored yet. We report the values in Table S1, along with a technical proof.

#### Mathematical details of the Hubbell regression

A generalized linear model (GLM; [39]) is a regression framework composed of three main elements: (i) an exponential dispersion family *f* (*y*_*i*_; *θ*_*i*_) which models the distribution of the outcome of interest *Y*_*i*_ in terms of a parameter *θ*_*i*_, independently for each *i* = 1, …, *N*; (ii) a linear predictor 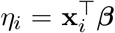, where **x**_*i*_ ∈ ℝ^*p*^ are covariates and ***β*** ∈ ℝ^*p*^ are the corresponding coefficients, and (iii) a link function *g* that expresses the relationship between *η*_*i*_ and the mean of *Y*_*i*_, so that **E**(*Y*_*i*_ | **x**_*i*_) = *µ*_*i*_ = *g*^−1^(*η*_*i*_). When *θ*_*i*_ = *η*_*i*_, the link is called *canonical*. GLMs enjoy several important statistical properties, including consistency, asymptotic unbiasedness [59], corrections for over-dispersion [60], and asymptotic efficiency [61].

The Hubbell regression is a GLM where the outcome of interest is the species richness observed in a community of size *n*_*i*_, denoted as 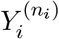, for the sampling sites *i* = 1, …, *N*. The exponential dispersion family that directs 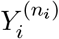 is

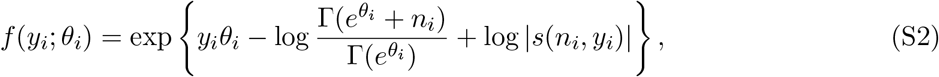

where *y*_*i*_ ∈ *{*1, …, *n*_*i*_*}*, and |*s*(*n, y*)| are the signless Stirling numbers of the first kind. Equation S2 is a re-arrangement of Equation 1 with 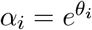. Following [56, 62], the mean and the variance functions for the distribution in Equation S2 are

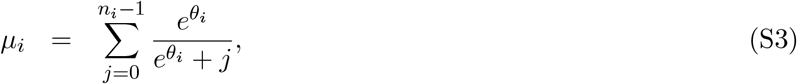

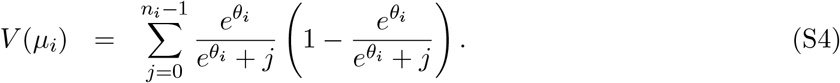

Both equations have an important ecological interpretation: Equation S3 is the expected species accumulation curve under Hubbell’s neutral theory, and Equation S4 defines its variance.

In this framework, the canonical GLM link is retained when *θ*_*i*_ = log *α*_*i*_ = *η*_*i*_ = **x**_*i*_^⊤^***β***. Hence, such a link establishes a log-linear model for *α*-diversity, allowing us to explicitly quantify how environmental covariates linearly scale the fundamental biodiversity of a site. In particular, a one-unit increase in the *r*th covariate *x*_*ir*_ translates into a 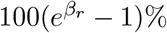 variation in *α*-diversity. While there is no analytical form for the link function *g*, the map between *µ*_*i*_ and *η*_*i*_ can be easily derived using root-finding algorithms during estimation.

#### Flexible link functions and regression-based diversity indices

In the canonical case, the rate at which *µ*_*i*_ grows with *n*_*i*_ is approximately logarithmic [63]. This motivates the asymptotic equivalence between Fisher’s *α* and the biodiversity parameter that governs the neutral theory. However, this assumption may be overly restrictive in practice, particularly for heavy-tailed abundance distributions, as observed empirically in arthropods [49]. To accommodate this, we adopt a general family of link functions *g* that grants flexible control of the growth rate of the accumulation curve [62].

The extended family of link functions we propose generalizes Equation S3 by introducing an additional parameter *σ* as follows:

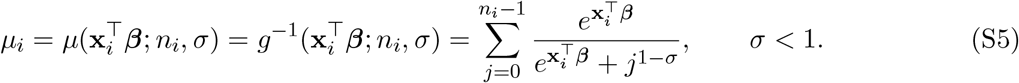

The above is a function of the linear predictor 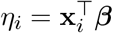 and depends both on the community size *n*_*i*_ and on *σ*. Extending the mean function as above allows us to generalize the logarithmic large-sample behavior of Equation S3. In particular, we prove that

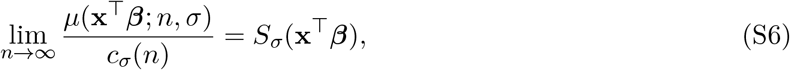

where *c*_*σ*_(*n*) is a function of both *n* and *σ* that captures the growth of *µ* with *n*. Specifically,

- (*Logarithmic growth*) - if *σ* = 0, then *c*_*σ*_(*n*) = log *n* and *S* 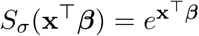;
- (*Polynomial growth*) - if *σ* ∈ (0, 1), then *c*_*σ*_(*n*) = *n*^*σ*^ and 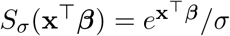;
- (*Finite richness*) - if *σ <* 0, then *c*_*σ*_(*n*) = 1 and 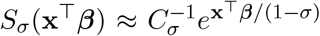, with *C*_*σ*_ a known constant.

Here, *S*_*σ*_(**x**^⊤^***β***) is the *regression-based diversity index*. When *σ* = 0, this quantity reduces to the canonical link, with 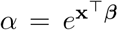. Since 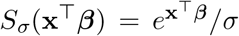 retains a log-linear structure, regression coefficients are interpreted in the same manner as the canonical link.

While flexible, the relationship between *n* and *Y* ^(*n*)^ in our model assumes a *common* growth rate for the accumulation curves across all sites. A common *σ* is what makes the regression-based diversity index *S*_*σ*_(*η*) and the regression coefficients comparable across sites, but this assumption may be restrictive in practice if the sampled population exhibits substantial heterogeneity or if there is evidence that environmental conditions affect the accumulation rate due to differences in sampling techniques. Solutions to address this limitation may involve additional hierarchical structures among sites and will be explored in future work.

#### Estimation

Because the link function *g* lacks a closed-form inverse, we invert it using a one-dimensional numerical root-finding algorithm (`uniroot` in R). In a saturated model, the regression-based diversity index *S*_*σ*_(*η*_*i*_) is obtained by solving the equation

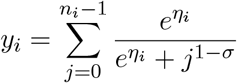

in terms of *η*_*i*_, for each *i* = 1, …, *N*. The resulting value for *S*_*σ*_(*η*_*i*_) when *σ* = 0 is close to Fisher’s *α* for sufficiently large *n*. As for the regression coefficients in a non-saturated model, estimation follows the well-known iteratively reweighted least squares algorithm [39]. The growth rate *σ* of the accumulation curve is learned in a data-driven manner via maximum likelihood.

In the general case, where *σ* is estimated from the data or chosen to be non-zero, Equation 2 and Equation 3 encode different growth rates with *n*. Therefore, we estimate *α* by first calculating *µ* = *g*^−1^(*η*) for a fixed *n*, and then solving Equation 2. Specifically, letting 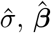, and 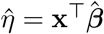 denote the maximum likelihood estimates for *σ*, ***β***, and *η*, we estimate the richness as 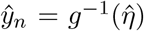 for any fixed community size *n*, and the associated Hubbell’s *α* by inverting

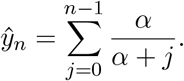

We call 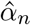 the solution to this procedure. It is also possible to show that

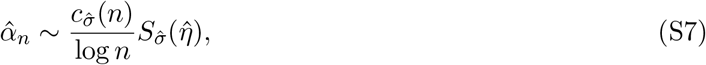

when *n* is sufficiently large. Note that Equation S7 holds with equality when *σ* = 0, the canonical link, in which case 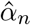 does not depend on *n*. The remaining biodiversity indices (Shannon, Simpson, Tsallis) are obtained by plugging 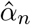 into their model-based formulas (Figure S7).

#### Jaccard-adjusted standard errors

In a GLM, observations are required to be conditionally independent. However, the same species, or BINs, may be detected across multiple sites, but counting the number of unique species ignores such information. Hence, it is necessary to account for the potentially ignored dependence during inference. Our solution is to treat the likelihood in the Hubbell regression as a composite marginal likelihood [64]. This is obtained as the product of individual likelihoods as if observations were independent, namely

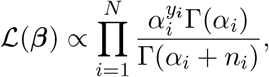

while correcting the standard errors of 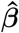 for heteroskedasticity.

The approach produces consistent estimates while mitigating the unexplained dependence during inference. We propose handling the relationship across sampling sites using spatial and heteroskedasticity-consistent standard errors [38], where spatial dependence is measured by the fraction of shared species between each pair of locations (Jaccard similarity index). Figure S3 depicts the relationship between the geographic distance between sampling points and this quantity. Calling **X** ∈ ℝ^*N ×p*^ the matrix of covariates, and **Ŵ** = diag*{ŵ*_1_, …, *ŵ*_*N*_ *}* the diagonal matrix of working weights, the variance-covariance matrix of the coefficients 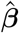 is estimated as

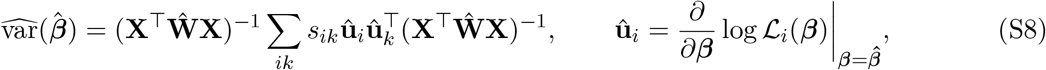

where *L*_*i*_(***β***) denotes the *i*th factor of the composite likelihood and *s*_*ik*_ ∈ [0, 1] represents a similarity index between sampling sites *i* and *k*. The standard errors are the square roots of the diagonal elements of 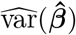. Exact comparisons between GLM standard errors based on the Fisher information, quasi-likelihood estimates, and the standard errors from Equation S8 are displayed in Figure S1.

#### The Global Malaise Trap Program

The Global Malaise Trap Program [19] is a large-scale sampling initiative aimed at assessing the global status of arthropod diversity; see http://globalmalaise.org/ and [65]. An interactive map of the current locations is available at the HubbellGLM Atlas at https://hubbellglm.org/. These data comprise 12,169,288 arthropod specimens collected using Townes-style Malaise traps deployed for about one week at each site (median 7 days). We sorted, photographed, and categorized the collected specimens morphologically to the closest known taxonomic rank, with 31.3% annotated to genus and only 13.0% to the species level. The DNA of each specimen was sequenced to amplify the cytochrome *c* oxidase I (COI) barcode region, and the resulting barcodes were uploaded to the Barcode of Life Data System and clustered into BINs; see [40] and [66]. We restrict our attention to the trapping events that captured at least 20 arthropods (*n* ≥ 20), and we discard samples in which the Malaise trap was damaged (e.g., by bears or extreme weather) or the collecting jar was found empty with no liquid. We further restrict the analysis to collections made up to 2024, the last year for which all environmental covariates are available, which excludes 535 events from 2025 and 2026. This resulted in 691,283 BINs and 11,523,563 specimens distributed across *N* = 15,176 collection events, gathered at 1,186 sites in 65 countries between 2007 and 2024. Sampling effort is spatially uneven: 6,260 collection events (41.2% of the total, and 48.8% of all specimens) come from 79 sites in Costa Rica, and a further 2,164 events (14.3%) from 619 sites elsewhere in North America. Coverage is also uneven in time: only 141 events (0.9%) predate 2012, and the length of the record varies sharply with latitude, from a median of 1 collection event per site in continental zones and 2 in polar ones to 13 in the tropics. Figure S8 shows when each site was sampled between 2007 and 2024, ordered by latitude. For each event, we observe the number of arthropods *n*_*i*_, the richness *y*_*i*_, and the fraction of shared BINs *s*_*ik*_ between every two events *i* and *k*.

#### Environmental covariates

We collect the following variables for both the sampling sites and a global raster on a regular 0.1° *×* 0.1° grid (1,508,557 land cells, ~11 km at the equator), for each year in the collection period (2007–2024). We use the latter for out-of-sample predictions. We obtained AET and PET values at each sample site from the TerraClimate dataset [67]. This indicator is derived from a one-dimensional water balance model, and it is reported monthly for any geographic location at a ~4 km spatial resolution. We annualized AET and PET by summing the monthly values at any latitude and longitude for the collection year, and report both in units of 10 mm/year. The realm of each site (divided into Afrotropic, Australasia, Indomalaya, Nearctic, Neotropic, or Palearctic) was downloaded from the WWF ecoregion data available in the Harvard Dataverse [68], while categorical Köppen–Geiger climatic zone classification (Continental, Dry, Polar, Temperate, and Tropical) was obtained using the kgc package [43]. For human influence, we use the HFP measure in [42]. This indicates the annual index of human pressure, ranging from 0 to 50, at ~1 km resolution. Annualized wetlands are downloaded from the Copernicus global land cover data [69]. Mean daily temperature at 2 meters, total precipitation, average relative humidity, average wind speed, and average vapor pressure deficit are downloaded from the ERA5 dataset [70] at ~10 km resolution. We use these data to compute weather anomalies for the collection window, as departures of the observed values from that cell’s own 1991–2020 day-of-year normal. Finally, we derive the area of the landmass on which each site lies and its great-circle distance to the nearest continental landmass from the Natural Earth 10m land polygons [71], treating a landmass as continental when its area exceeds 5 *×* 10^6^ km^2^.

#### Modeling the influence of evapotranspiration and human footprint on diversity

To test the joint effect of climatic factors and human footprint on biodiversity, we estimate a sequence of nested models of increasing complexity. Each model below adds the described covariates to the preceding one.

- **M0** Null model. A model with an intercept only. This model captures the dependence between the community size *n* and the richness *y* through the accumulation curve alone, without covariates. If *σ* = 0, this automatically entails a logarithmic growth.
- **M1** Evapotranspiration. A simple GLM with AET as the sole covariate. This tests the univariate relationship between diversity and its putative primary driver.
- **M2** Realm fixed effects. We further add a fixed effect to control for the realms. This captures the geographic area of each sampling site at the macro level.
- **M3** Seasonality. To account for variations in diversity that are uniquely driven by the time of collection, we add the collection week to the model using a one-degree Fourier transform. That is, we include both 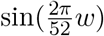 and 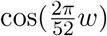, where *w* is the ISO week of collection in the year, restricted to be between 1 and 52. If the site is below the equator (latitude *<* 0), we shift *w* by 26 weeks, so that *w* = 26 represents the mid-point of the year everywhere.
- **M4** Latitude effect. We control for the residual spatial variation in biodiversity that is not explained by AET in a nonparametric way, modeling the latitude of each sampling site via a natural cubic spline with 6 degrees of freedom. We found such a choice to be the best compromise between flexibility and fit.
- **M5** Human footprint and climatic zone. We assess the effect of the human footprint (HFP) and its interaction with the Köppen–Geiger climatic zones. This captures the interplay between human activity and climate in an interpretable manner.
- **M6** Climatic variables and additional controls. We further include climatic and geographic factors in the model, specifically annual average wind speed, wetlands, and the residuals from regressing mean annual temperature on latitude and latitude squared. We also control for the number of collection days, which is 7 under normal conditions, and include a year trend. To control for island versus mainland effects, we also include log_10_ of the area (in km^2^) of the landmass on which each site lies, and log_10_ of one plus its great-circle distance (in km) to the nearest continental landmass.
- **M7** Weather variations. We finally control for short-term weather fluctuations during the collection window by adding four anomalies: the departures of the observed temperature, wind speed, total precipitation, and relative humidity from that cell’s own 1991–2020 day-of-year normal, averaged over the days the trap was deployed.

These models capture the main effect of AET and the interaction between climatic areas and HFP. For models M0 to M7, we estimate a Hubbell regression with a canonical link (*σ* = 0) and one in which *σ* is fixed across all specifications to its maximum likelihood estimate under the full model M7 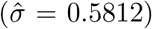, so that regression coefficients are directly comparable across the nested sequence. Regression coefficients and standard errors are reported in the Supplementary material in Table S3 and Table S4.

#### Benchmarking Hubbell regression with other models

To evaluate Hubbell regression against alternatives, we compare it with seven estimators fitted on identical training rows and scored on identical held-out rows. Our benchmarks are 1) a Poisson regression model for *y*_*i*_−1 with a log-link; 2) a negative binomial regression for *y*_*i*_ −1 with a log-link; 3) a linear model for log *y*_*i*_, with ns(`n_i`, `df` = 10) as a flexible covariate; 4) a linear model for log 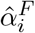, where 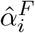 is Fisher’s *α* for sample *i*; and 5) the Hubbell regression under the canonical link (*σ* = 0), which isolates the contribution of the polynomial link with the mean structure held fixed. To ensure a fair comparison of species accumulation rates, we consider two variants of the Poisson and negative binomial models. The first uses log *n*_*i*_ as a further covariate, so as to mimic the polynomial relationship between *n*_*i*_ and *y*_*i*_ in the Hubbell regression with link as in Equation S5. The second uses an offset equal to log(*γ* + log *n*_*i*_) instead, mirroring the logarithmic growth of the canonical Hubbell link, where *γ* ≈ 0.5772 is the Euler–Mascheroni constant. These choices are motivated by Theorem S2, which illustrates an asymptotic equivalence between the canonical Hubbell regression and a Poisson regression with offset, under large *n*.

We evaluated all estimators under specification M7 and four cross-validation partitions: (i) 10 random folds over collection events; (ii) 10 folds formed by assigning 500 km spatial blocks at random; (iii) 10 folds formed by *k*-means clustering of site coordinates; and (iv) leave-one-realm-out, holding out each of the six biogeographic realms in turn. The two spatial partitions are computed with the `blockCV` package [72] on Mollweide equal-area coordinates (ESRI:54009), and in partitions (ii)–(iv) all events from a site share a fold, so that a held-out fold is an unsampled region. Partitions (i)–(iii) are repeated over 10 random replicates, giving 100 folds each, while (iv) is deterministic and yields 6 folds. Under partitions (iii) and (iv) the realm term is removed from every estimator, since a held-out realm’s level cannot be estimated from training data that never contain it. Figure S9 displays four examples of the partitions in scenarios (i)–(iv).

Within each fold, *σ* is re-estimated as described in Algorithm S1. For each fitted model, we calculate the root-mean-squared error (RMSE) between *y*_*i*_ and its predicted value, both in-sample and in the held-out folds. For every fold, we calculate the paired percentage difference between each competitor and the Hubbell regression with polynomial growth, obtained as 100 *×* (RMSE_comp_ − RMSE_Hub_)*/*RMSE_Hub_, where RMSE_comp_ is the RMSE from a competitor model, and RMSE_Hub_ is the RMSE from the Hubbell regression with polynomial link, for each fold and replicate. Figure S5 shows the results.

We find that Hubbell regression attains the lowest in-sample error against every alternative, in every fold of every partition, and the lowest out-of-sample error in almost all scenarios (Figure S5). The contrast is sharpest against the logarithmic link: fixing *σ* = 0 raises the out-of-sample RMSE by 34.5% to 43.7% depending on the partition. Against the Poisson, negative binomial, and log-richness regressions, the out-of-sample gap narrows as the partitions become more separated, from 4.3–17.5% under random folds to parity under *k*-means and leave-one-realm-out, even though those models fit the training data worse. However, Hubbell regression is the only model that yields an admissible accumulation curve (i.e., one where *y* ≤ *n* everywhere; see Figure S6). This confirms the inherent advantages of our framework for modeling richness over existing alternatives when using the same environmental covariates.

#### Global predictions and scenario evaluation

We obtain the results in Figure 3 as follows. We predict richness on the 0.1° global grid for the reference year 2024, fixing the community size to *n* = 2000 individuals, the collection week to *w* = 26, the collection duration to 7 days, and all weather anomalies to zero, so that maps differ only through the environmental covariates. Each scenario compares a baseline prediction *µ*_0_ to a counterfactual *µ*_1_ in which one covariate is modified (HFP set to zero, HFP raised by 5 units, or AET raised by 10% of its local value), and reports the relative change *R* = 100(*µ*_1_*/µ*_0_ − 1) at any point along the accumulation curve. Confidence intervals for *R* are obtained via parametric bootstrap, drawing ***β*** from a Gaussian distribution centered at 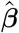 with the Jaccard-adjusted covariance matrix in Equation S8.

Because a fitted model should not be read outside the environmental range it was estimated on, we compute the multivariate environmental similarity surface [MESS; [73]] for every grid cell, within its own climatic zone, over the continuous predictors of the model: AET, latitude, HFP, wind speed, wetlands, the residual temperature term, and the two island terms on the log_10_ scale used in the regression. Negative MESS values indicate cells outside the training envelope. We mask cells with MESS *<* −25, which removes 8.0% of land; masking is negligible in temperate (0.9%), dry (1.5%), and tropical (2.1%) zones but substantial in polar regions (39.6%), where the GMTP contains few sites. The MESS for year 2024 is reported in Figure S10. All scenario statistics reported in the main text are computed on the unmasked cells.

#### Variations in AET and HFP for additional biodiversity indices

To show how the estimated interplay between AET and HFP can be translated to other commonly used biodiversity indices, we calculate our predictions for the richness *ŷ*_*n*_ using the link function in Equation 3. We retrieve the associated 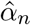 as the solution of Equation 2 when *µ* = *ŷ*_*n*_. The remaining indices are simple functions of *α* (Table S1), and can be visualized for, e.g., a fixed community size *n* = 2000. Consistent with our regression coefficient estimates, we find that higher AET is associated with higher Shannon diversity and with a lower Simpson index. Note that the model-based Simpson index 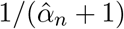 measures the probability that two individuals drawn from the sample belong to the same species (Table S1), so a lower value corresponds to a more diverse community, and the two indices therefore move consistently. Instead, higher HFP levels are associated with lower values in dry and tropical regions, thereby flattening the diversity curves, whereas polar regions exhibit the opposite trend (Figure S7). Figure S11 translates the predictions for all biodiversity indices at the global scale.

#### The HubbellGLM package

All models in this paper are estimated via the R package `HubbellGLM`. To facilitate usage, we designed it to mimic the R syntax of the glm function, that is

`HubbellGLM(cbind(n, y) ~., family = hubbell(sigma = 0), data = data)`.

The output is compatible with many base R functions for model evaluation, such as `summary, plot, predict, residuals, deviance, step`, and more. By default, we keep *σ* fixed to a pre-specified value, and estimate it separately via the function `estimate_sigma(formula, data)`. The variance-covariance matrix for shared-species standard error adjustment is coded via the function `vcov_shared()`, and the output can be used in combination with the `coeftest` function of the package `lmtest`.

### Supplementary text

#### Algorithm S1

**Iteratively Reweighted Least Squares for the Hubbell regression Input:** design matrix **X**, species richness *y*_*i*_, community size *n*_*i*_, initial ***β***^(0)^, fixed value for *σ <* 1.

**For** *t* = 0, 1, 2, … until convergence:

• **1. Update the mean**

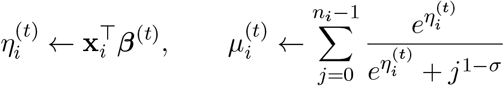
• **2. Retrieve diversity** *α*_*i*_ **and variances** By using root-finding:

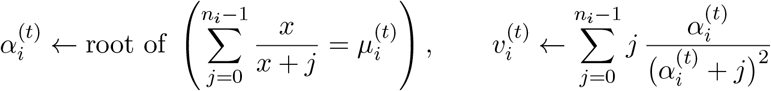
• **3. Compute working weights**

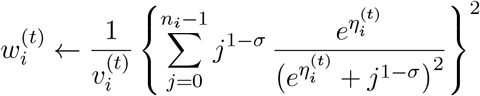
• **4. Update regression coefficients**

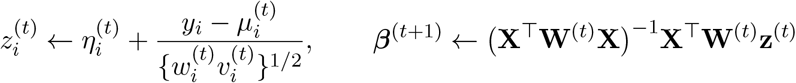
**5. Profile optimization of** *σ* **(Optional)** Let 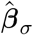 denote the converged solution of the above IRLS algorithm for a conditionally fixed *σ*, and 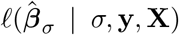 denote the log-likelihood at that point. Estimate the optimal accumulation rate 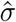 by maximizing the profile log-likelihood via bounded quasi-Newton optimization (e.g., utilizing `nlminb` in `R`):

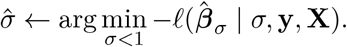

The pair 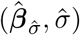 jointly minimizes −ℓ(***β***_*σ*_ | *σ*, **y, X**).

#### Technical proofs

##### Proposition S1

*Let π*_1_ ~ Beta(1, *α*) *for α >* 0, *and call* **E**(*I*_*f*_) = **E**(*f* (*π*_1_)*/π*_1_). *Then, the following holds*.

1. *Shannon index: when f* (*x*) = −*x* log *x, then* **E**(*I*_*f*_) = *ψ*(*α*+1)−*ψ*(1), *where ψ*(*x*) *is the digamma function*.
2. *Simpson index: when f* (*x*) = *x*^2^, *then* **E**(*I*_*f*_) = 1*/*(*α* + 1).
3. *Tsallis diversity: when* 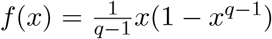 *for any q >* 0 *with q* ≠ 1, *then* 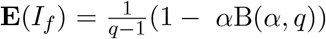, *where* B(*a, b*) *is the beta function*.

*Proof*. The statement follows from properties of the beta distribution, whose probability density function for *x* ~ Beta(1, *α*) is *p*(*x*) = *α*(1 − *x*)^*α*−1^.

1. Shannon index:

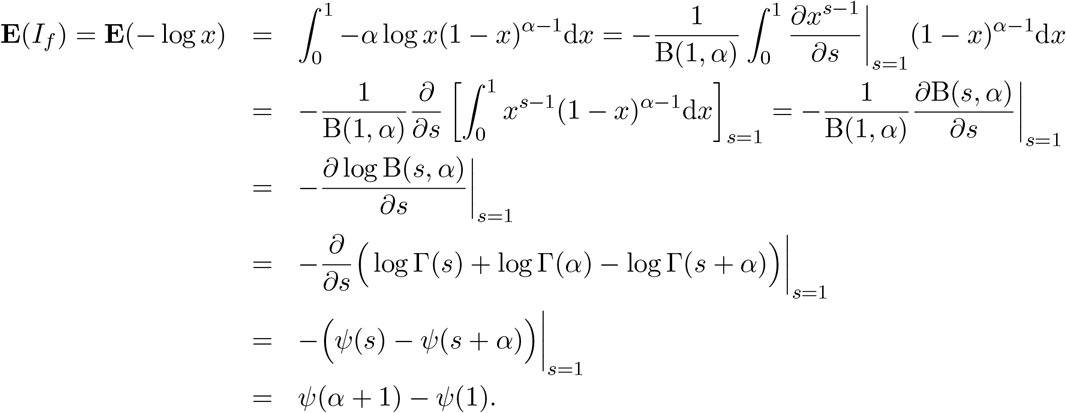
2. Simpson index: 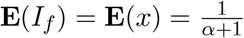, by definition of expected value of the beta distribution.
3. Tsallis diversity: 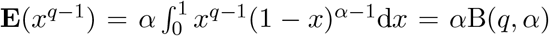, by definition of beta function. Moreover, since the beta function is symmetric, B(*q, α*) = B(*α, q*). The proof is completed using the linearity of expectations, since 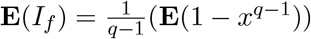.

##### Theorem S1

*Let* 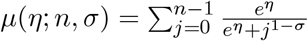, *for σ <* 1 *and η* ∈ ℝ. *Then, it holds that*

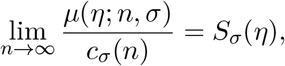

*where:*

1. *If σ* = 0, *then c*_*σ*_(*n*) = log *n and S*_*σ*_(*η*) = *e*^*η*^
2. *If σ* ∈ (0, 1), *then c*_*σ*_(*n*) = *n*^*σ*^ *and S*_*σ*_(*η*) = *e*^*η*^*/σ*
3. *If σ <* 0, *then c*_*σ*_(*n*) = 1 *and* 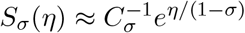, *where C*_*σ*_ *is a constant*.

*Proof*. Consider the function 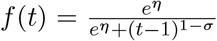, for *η* ∈ ℝ, *σ <* 1, and any *t* ≥ 1. Then, we have that 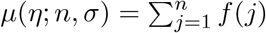. Since *f* (*t*) is monotonically non-increasing in *t*, it holds that

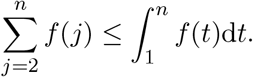

Adding *f* (1) = 1 to both sides of the inequality above then yields

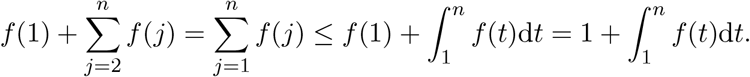

Moreover, by monotonicity of *f* (*t*) it also holds that

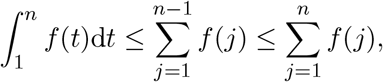

where the rightmost inequality is always true because *f* (*t*) ≥ 0 for any *t*. Combining both sides of the inequalities and dividing by *c*_*σ*_(*n*) leads to the following bounds, which hold for every *n* ≥ 2:

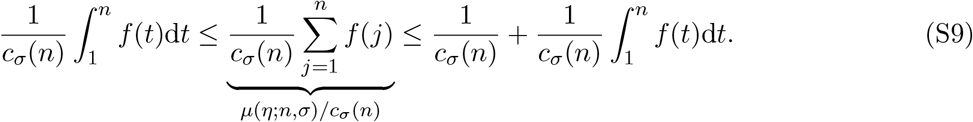

The rest of the proof follows from evaluating the integral and taking the limit as *n* → ∞. In particular,

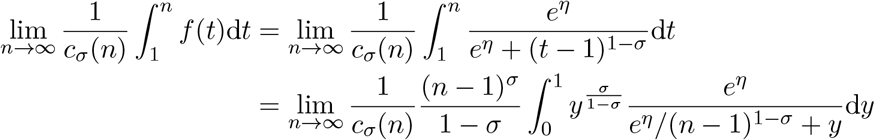

When *σ* = 0, then *c*_*σ*_(*n*) = log *n* and

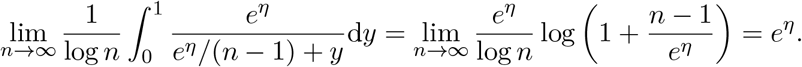

When *σ* ∈ (0, 1), then *c*_*σ*_(*n*) = *n*^*σ*^ and

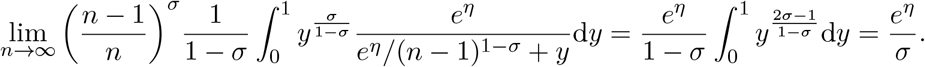

Since in both cases 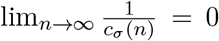, the lower and upper bounds in Equation (S9) converge to the same limit as *n* → ∞, and so does *µ*(*η*; *n, σ*)*/c*_*σ*_(*n*), which lies between them for every *n*. This concludes the proof for parts 1 and 2. As for part 3, when *σ <* 0 we have *c*_*σ*_(*n*) = 1, and it holds that

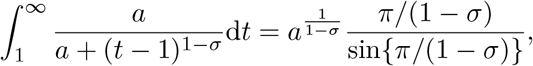

for any *a >* 0 [62]. Moreover, since *f* (*t*) ≥ 0, the sequence *µ*(*η*; *n, σ*) is non-decreasing in *n* and, by Equation (S9), bounded, so that its limit exists. Hence, since Equation (S9) holds for every *n*, taking the limit as *n* → ∞ on both sides leads to

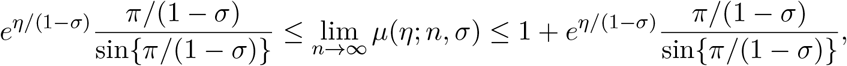

which completes the proof, with *C*_*σ*_ = sin*{π/*(1 − *σ*)*}/{π/*(1 − *σ*)*}*.

##### Theorem S2

*Call γ* = 0.5772 … *the Euler–Mascheroni constant and let σ* = 0 *in the Hubbell regression model. If n*_*i*_ → ∞ *for observation i, then we have*

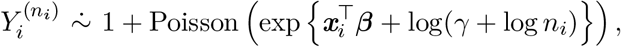

*where* 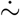 *denotes approximation in distribution*.

*Proof*. The proof proceeds as follows. Recall that the canonical link implies that

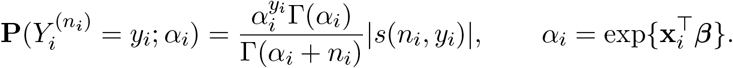

Let *a*(*n*) ~ *b*(*n*) indicate that lim_*n*→∞_ *a*(*n*)*/b*(*n*) = 1. From [74], Section 24.1.3, we have the following asymptotic approximation of the signless Stirling numbers of the first kind:

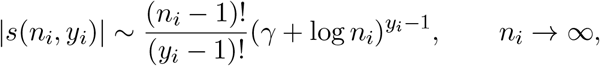

where *γ* = 0.5772 … is the Euler–Mascheroni constant. Hence, plugging in the above approximations yields

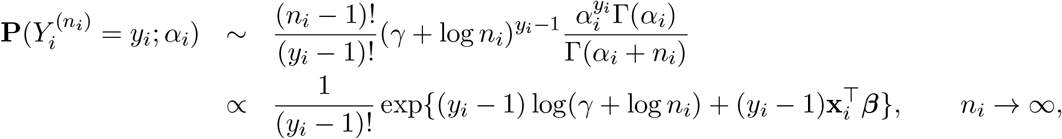

where the proportionality is taken with respect to *y*_*i*_. The above distribution is the kernel of a 1-shifted Poisson distribution with mean 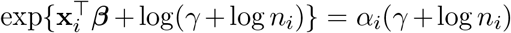. This completes the proof.

**Figure S1.**
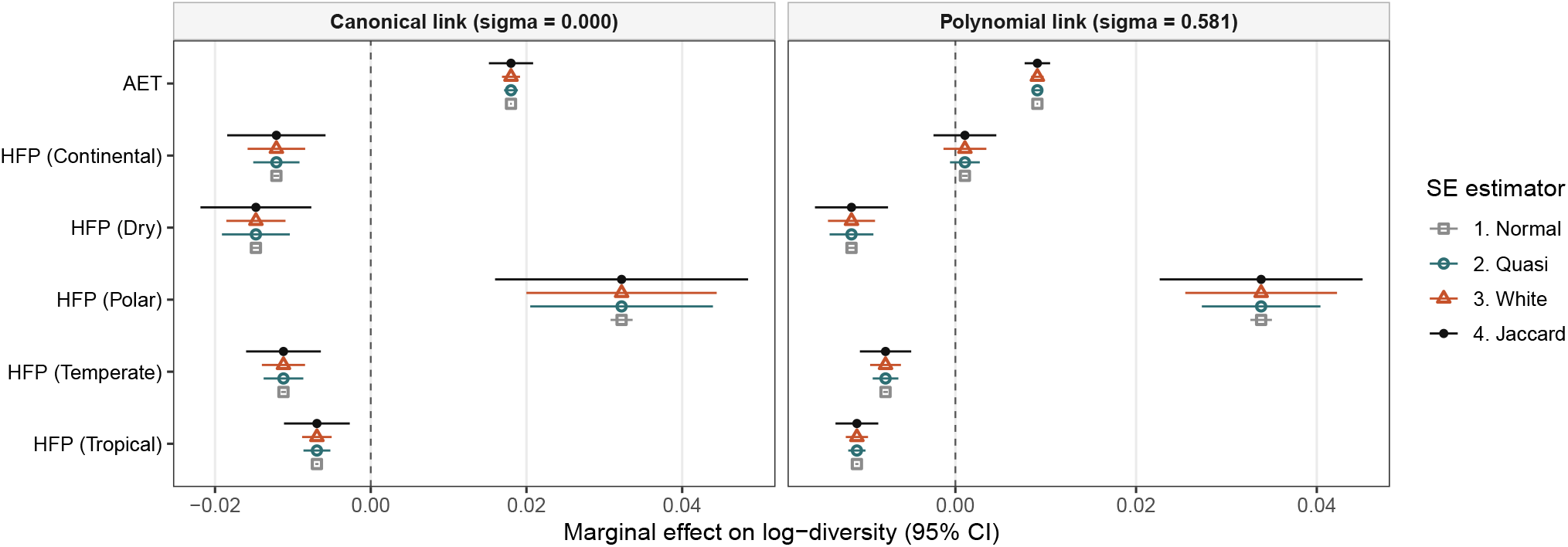
Comparison of 95% confidence interval ranges for AET and HFP across link functions and standard error estimators. Normal indicates the standard errors calculated using default glm routines, while quasi uses the quasi-likelihood approach for the Hubbell family. White are White heteroskedasticity-consistent standard errors, and Jaccard are adjusted for shared species.

**Figure S2.**
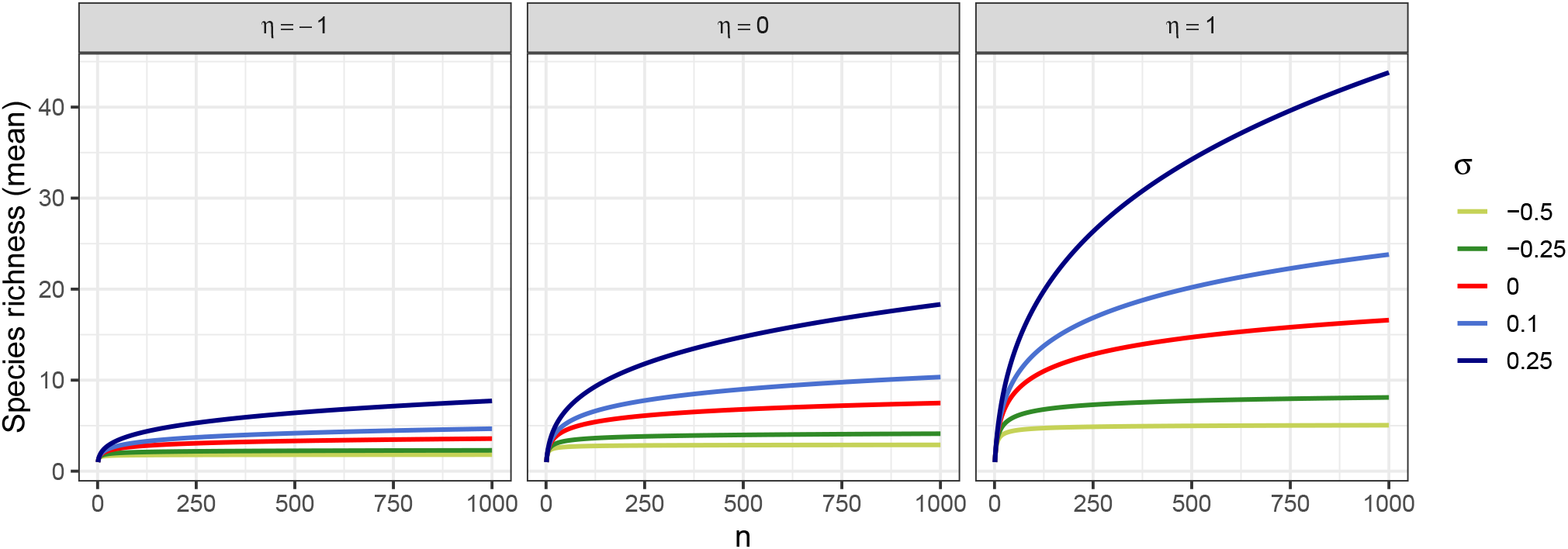
Examples of accumulation curves for the polynomial link function. When *σ* = 0, the curve grows logarithmically. Values for *σ <* 0 lead to a finite asymptote, while *σ* ∈ (0, 1) leads to polynomial growth.

**Figure S3.**
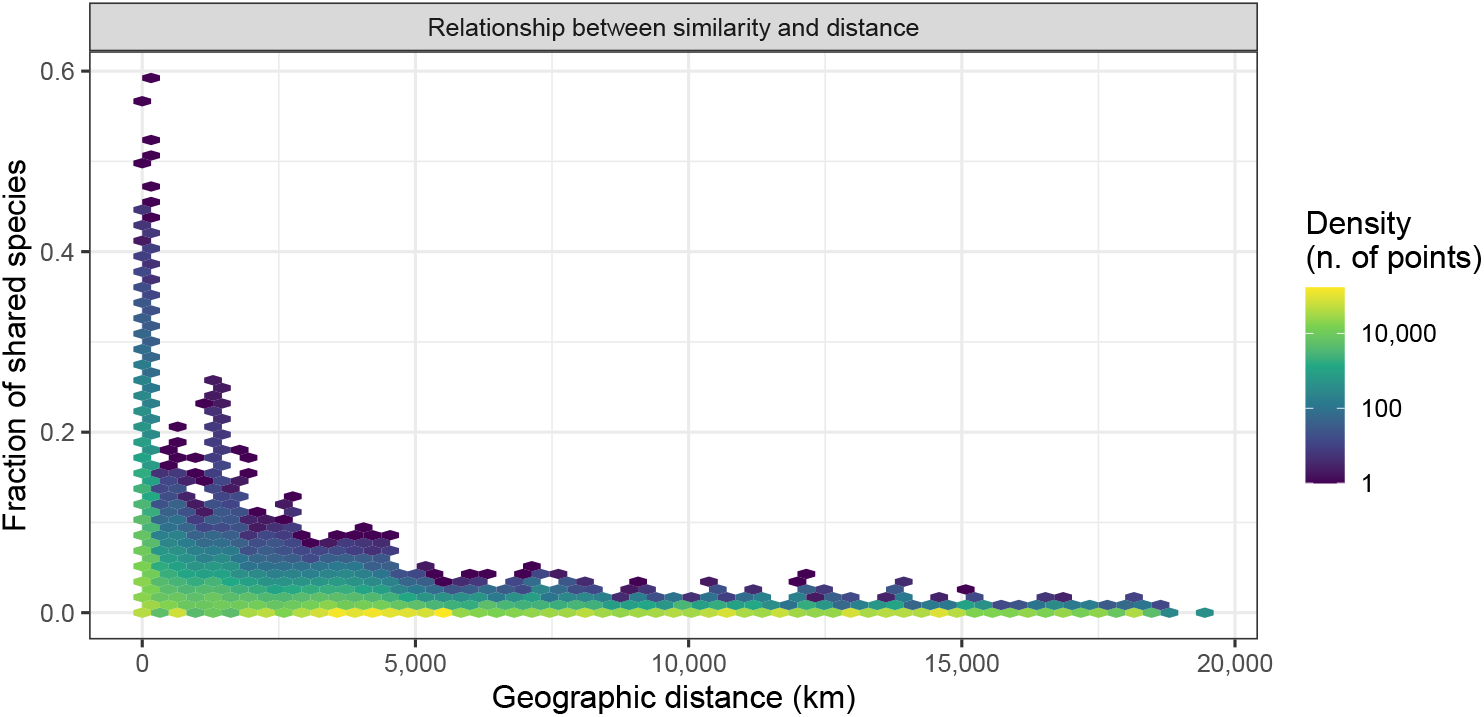
Relationship between the geographic distance between each collection event and the Jaccard similarity index. Color intensity indicates the number of points that fall within each bin.

**Table S1.** Overview of diversity indices, their interpretations, and formulas. Here, *y* indicates the sample richness, that is, the total number of unique species observed. In addition, *p*_*k*_ represents the relative frequency of the *k*th species in the sample. “Model-based” refers to the modeling setup of the unified neutral theory of biodiversity and biogeography. Note that *ψ*(*x*) = Γ^*′*^(*x*)*/*Γ(*x*) is the digamma function, and B(*a, b*) is the beta function.

| Diversity index | Interpretation | Sample-based | Model-based |
| --- | --- | --- | --- |
| α-diversity | Effort-independent measure of local richness | $y = \hat{\alpha}^F \log(1 + n/\hat{\alpha}^F)$ | $y = \sum_{j=0}^{n-1} \frac{\alpha}{\alpha+j}$ |
| Shannon | Entropy in the sample composition | $-\sum_{k=1}^y p_k \log p_k$ | $\psi(\alpha+1) - \psi(1)$ |
| Simpson | Probability that two individuals randomly drawn from a sample are from the same species | $\sum_{k=1}^y p_k^2$ | $1/(\alpha+1)$ |
| Tsallis | Generalized notion of entropy, encompassing various indices based on $q > 0, q \neq 1$ | $\frac{1}{q-1}(1 - \sum_{k=1}^y p_k^q)$ | $\frac{1}{q-1}(1 - \alpha B(\alpha, q))$ |

**Figure S4.**
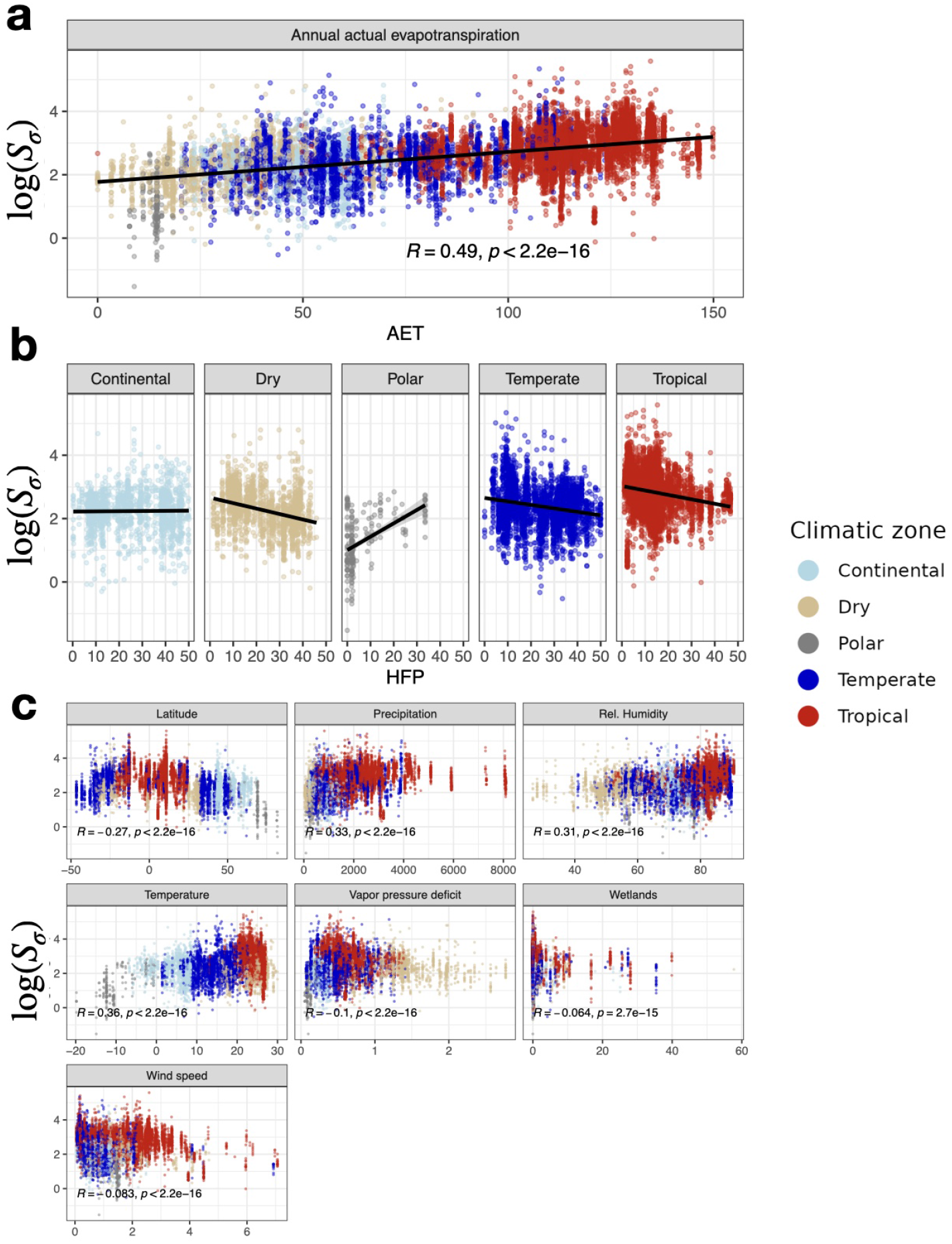
Relationship between environmental covariates and diversity. Colors in the plot denote climatic zones. (**a**) Scatterplot of annual actual evapotranspiration (AET) and the logarithm of the regression-based diversity (log *S*_*σ*_), calculated in each sample under a saturated model with *σ* = 0.5812. See Materials and Methods for exact calculation. (**b**) Scatterplot between human footprint (HFP) and log *S*_*σ*_, by climatic zone. (**c**) Scatterplot of annual covariates and log *S*_*σ*_. In each plot, *R* indicates the correlation coefficient, and *P* the associated *P* value to test whether it is significantly different from zero.

**Figure S5.**
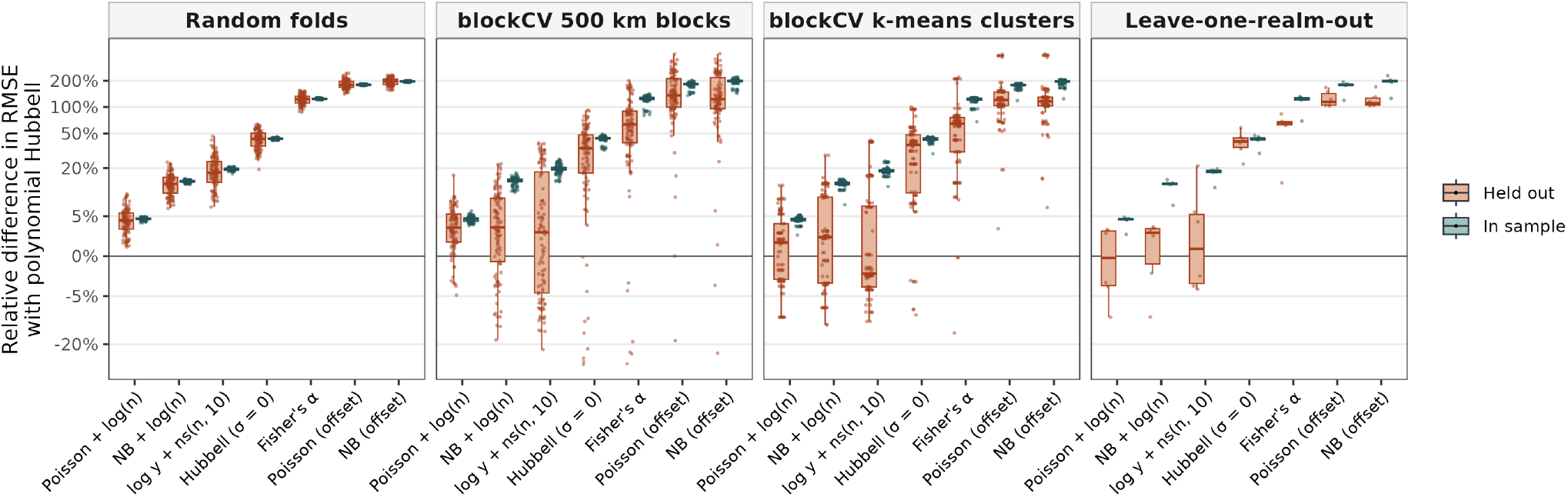
Benchmarking simulation via cross-validation. Boxplots depict the in- and out-of-sample % difference in root-mean-squared error (RMSE) of each competitor model against the Hubbell regression with polynomial growth, one point per fold, under specification M7. Values above zero indicate that the competitor predicts the observed richness less accurately. The three randomized strategies use 10 folds over 10 replicates, while leave-one-realm-out is deterministic and yields only 6 folds.

**Figure S6.**
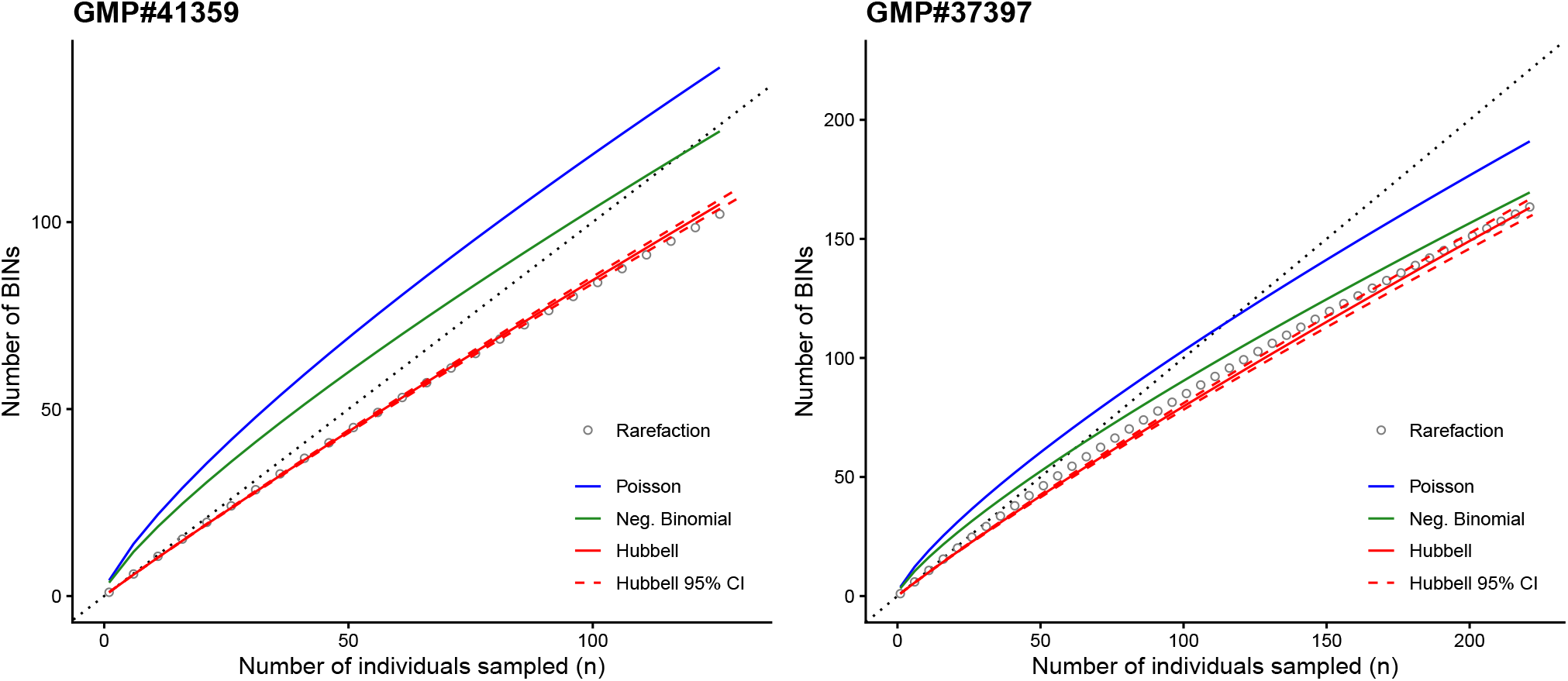
Observed rarefaction and predicted accumulation curves. Collection events GMP#41359 (Peru, left) and GMP#37397 (Thailand, right). The dotted line is the 45-degree line where *y* = *n*, the maximum possible richness. Poisson and negative binomial regressions predict more BINs than individuals sampled over part or all of the curve in these two examples.

**Figure S7.**
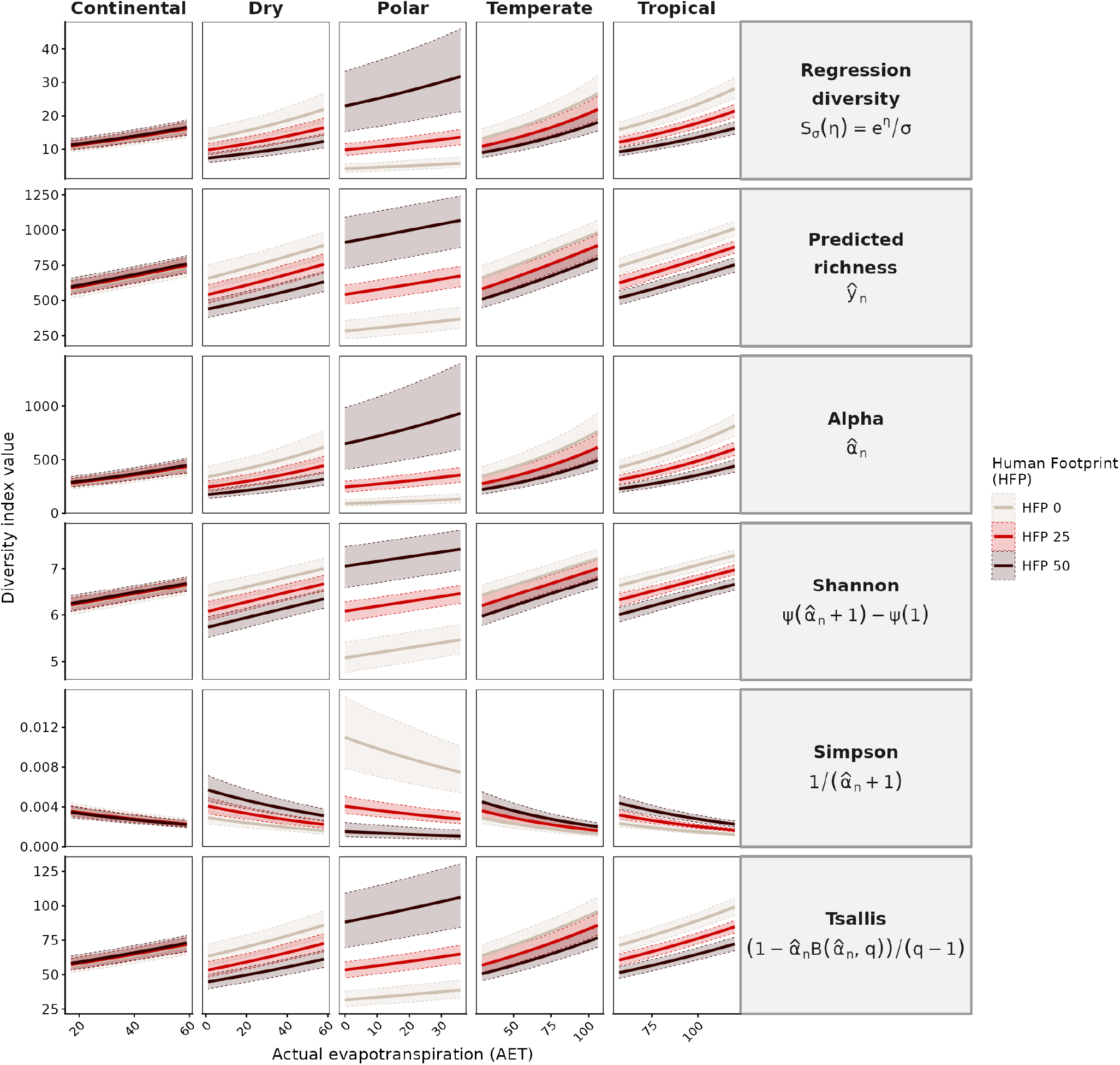
Variations of AET and HFP across climatic zones and biodiversity indices. All curves are predictions from model M7 for the year 2024, fixing the community size to *n* = 2000, *σ* = 0.5812, the collection week to the 26th week of the calendar year, and the collection duration to 7 days. Covariates are set as in Figure 2, and the realm is the most common realm of each zone on the global grid (Nearctic for Polar, Palearctic for Continental, Dry, and Temperate, Neotropic for Tropical). Here 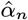 corresponds to the solution of Equation 2 when *n* = 2000 and *µ* = *ŷ*_*n*_. Bands around the lines indicate 95% bootstrap confidence intervals, computed using Jaccard-adjusted standard errors. Ranges on the x-axis correspond to plausible AET ranges in each zone at a global scale.

**Figure S8.**
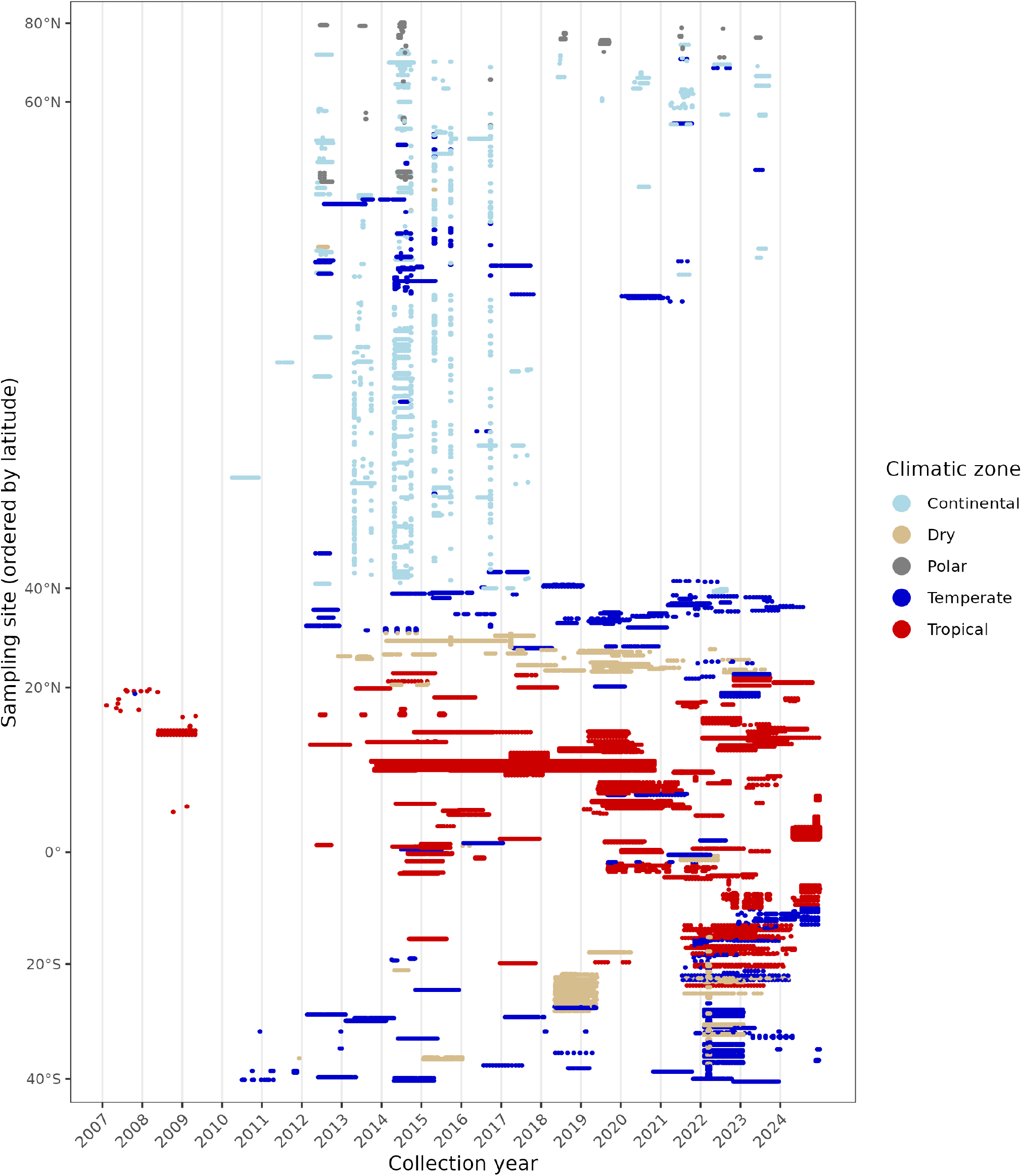
Collection history of each sampling site in the GMTP, ordered by latitude. Colors in the plot denote climatic zones. Each point marks a one-week collection event at a given site, positioned by its calendar date.

**Figure S9.**
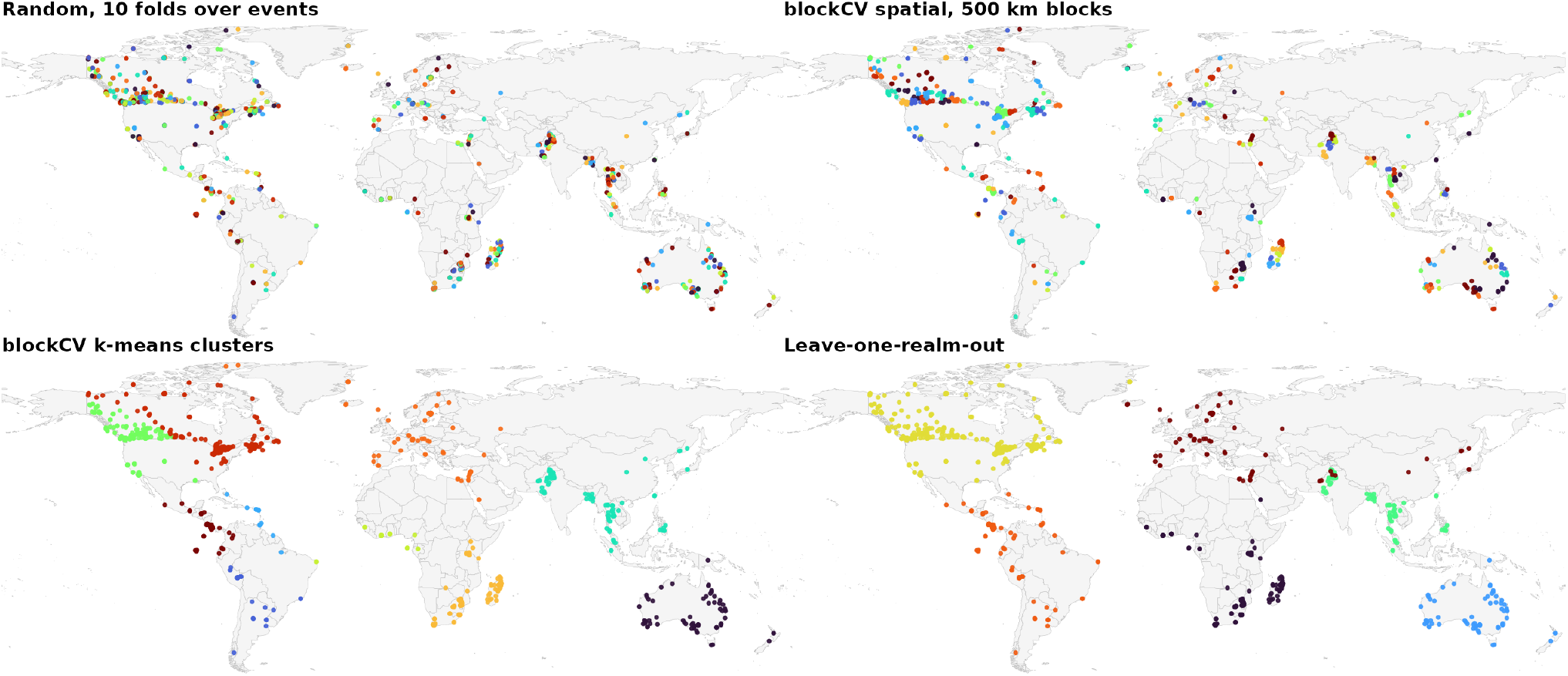
Example of partitions in the simulation study. Different colors indicate sites in different folds.

**Figure S10.**
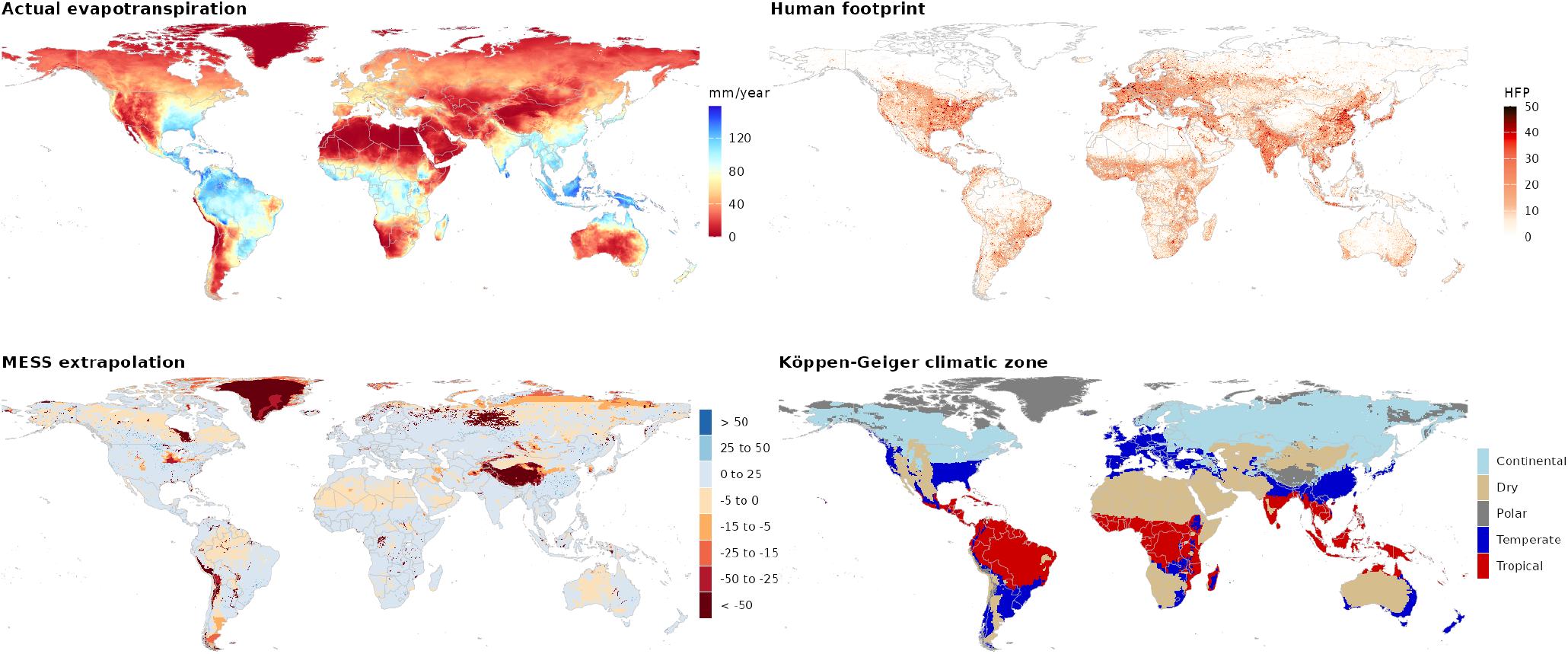
Global distribution of main covariates under analysis and MESS extrapolation score. Values refer to the year 2024, which is the most frequent in the collection.

**Figure S11.**
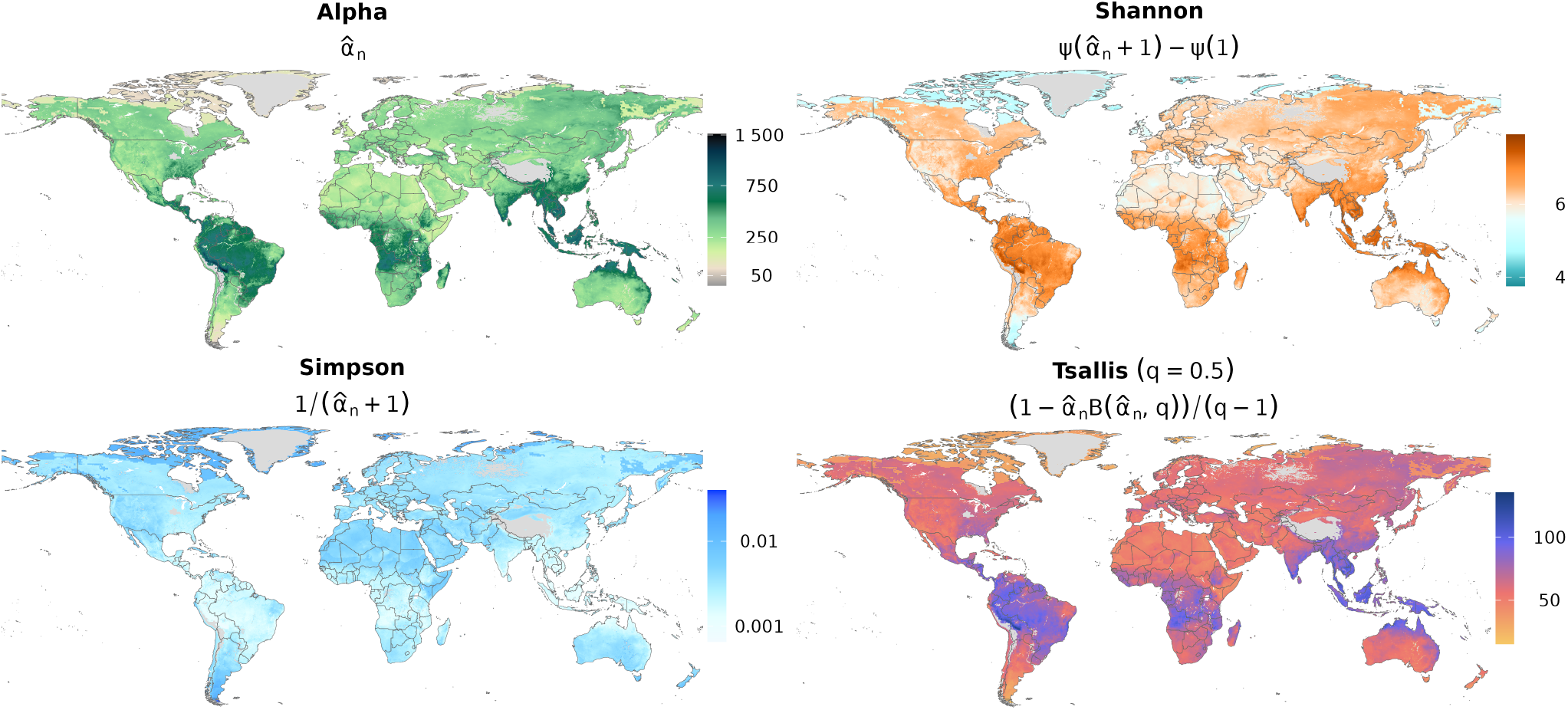
Global values for each biodiversity index in the year 2024. The symbols *ψ*(*x*) and B(*a, b*) denote the digamma function and the beta function, respectively.

**Table S2.** Marginal explanatory power (Pseudo-*R*^2^) across model specifications, with *σ* = 0.608. As in Table S3, *σ* is held fixed across all specifications at its maximum likelihood value under the most complex model. Here, however, this reference model excludes AET and the other candidate drivers compared in the table, so that the choice of *σ* does not favor any of them; this is why *σ* differs slightly from the value 0.5812 used elsewhere. The baseline deviance for each specification is reported below its name. The marginal drop in deviance for each predictor is reported below the *R*^2^ in parentheses. Empty cells indicate variables that were not statistically significant (*P >* 0.05, Chi-square test with Jaccard-adjusted *P* values). The highest *R*^2^ and largest deviance drop per specification are highlighted in bold. The column Form indicates the test function form for the covariate (linear or natural cubic spline). **AET**: annual actual evapotranspiration; **PET**: annual potential evapotranspiration; **VPD**: annual mean vapor pressure deficit; **Temp**: annual mean temperature; **Precip**: annual total precipitation; **Humid**: annual mean relative humidity. *n* = 15,176 collection events.

| Driver | Form | 1 | + Realm | + Week | + ns(Lat, 6) | + Zone×HFP | + Controls | + Anomalies |
| --- | --- | --- | --- | --- | --- | --- | --- | --- |
|  |  | (1,065,442) | (850,507) | (845,919) | (756,634) | (675,007) | (512,727) | (509,315) |
| **AET** | Linear | 0.305 | 0.321 | 0.326 | 0.376 | 0.402 | 0.552 | 0.556 |
|  |  | (325,032) | (127,299) | (127,759) | (91,527) | (37,743) | (35,204) | (36,424) |
|  | Spline | **0.309** | **0.327** | **0.332** | 0.378 | 0.406 | **0.555** | **0.559** |
|  |  | **(329,504)** | **(133,479)** | **(133,870)** | (93,626) | (41,649) | **(38,132)** | **(38,937)** |
| **PET** | Linear | 0.059 | 0.205 | 0.209 | 0.329 | 0.397 | 0.534 | 0.538 |
|  |  | (63,205) | (3,511) | (3,107) | (41,600) | (32,352) | (16,400) | (17,267) |
|  | Spline | 0.163 | 0.237 | 0.243 | 0.347 | 0.409 | 0.536 | 0.540 |
|  |  | (173,524) | (37,755) | (39,031) | (60,668) | (45,336) | (18,361) | (19,151) |
| **VPD** | Linear | 0.044 | 0.257 | 0.260 | 0.365 | 0.411 | 0.534 | 0.538 |
|  |  | (46,695) | (58,551) | (56,970) | (80,502) | (47,373) | (16,071) | (17,050) |
|  | Spline | 0.104 | 0.279 | 0.281 | 0.377 | 0.424 | 0.537 | 0.542 |
|  |  | (111,320) | (81,872) | (80,223) | (92,788) | (61,806) | (19,642) | (20,928) |
| **Temp** | Linear | 0.106 |  |  | 0.305 | 0.397 | 0.520 | 0.523 |
|  |  | (112,876) |  |  | (16,200) | (32,397) | (977) | (1,018) |
|  | Spline | 0.235 | 0.281 | 0.287 | 0.341 | **0.426** | 0.539 | 0.543 |
|  |  | (250,164) | (84,769) | (86,660) | (54,694) | **(63,100)** | (21,504) | (22,047) |
| **Precip** | Linear | 0.112 | 0.214 | 0.220 | 0.299 | 0.370 | 0.524 | 0.527 |
|  |  | (119,589) | (13,498) | (14,364) | (9,861) | (3,860) | (5,932) | (5,723) |
|  | Spline | 0.185 | 0.243 | 0.247 | 0.323 | 0.380 | 0.528 | 0.532 |
|  |  | (197,637) | (43,458) | (44,073) | (34,811) | (14,298) | (10,031) | (10,370) |
| **Humid** | Linear | 0.175 | 0.255 | 0.259 | 0.351 | 0.390 | 0.531 | 0.535 |
|  |  | (186,569) | (57,245) | (56,651) | (65,485) | (25,122) | (13,014) | (13,869) |
|  | Spline | 0.235 | 0.290 | 0.295 | **0.381** | 0.425 | 0.534 | 0.538 |
|  |  | (250,214) | (94,200) | (94,566) | **(96,743)** | (61,944) | (15,933) | (16,716) |

**Table S3.**
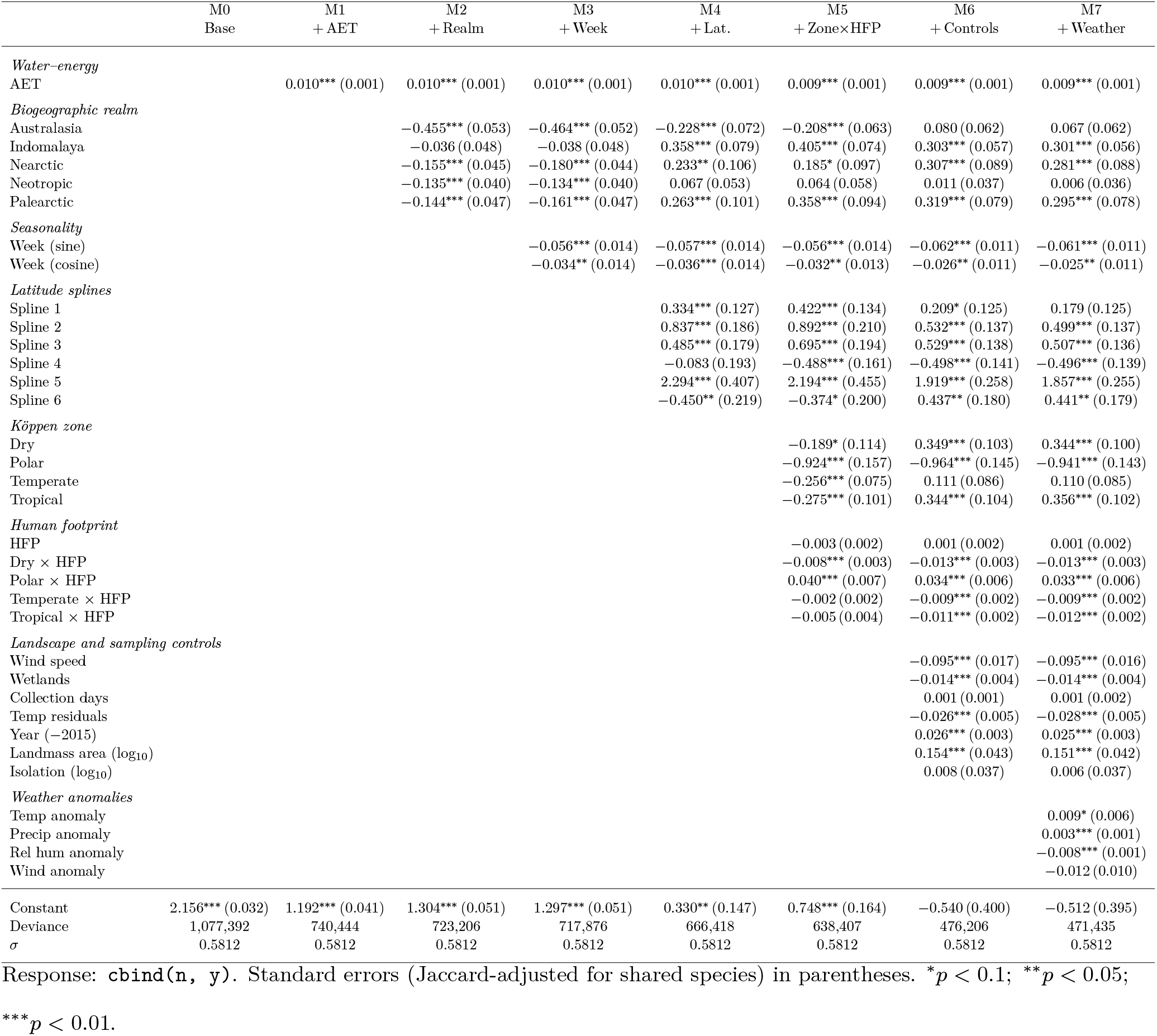
Stepwise Hubbell regression estimates (polynomial link). Coefficients are added in nested blocks from the base intercept-only model (M0) to the full weather-anomaly model (M7). Realm and climate-zone coefficients are relative to the omitted reference category. The accumulation-rate parameter *σ* is held fixed at its maximum likelihood value in the full specification (M7) for every model, so that coefficients are directly comparable across the nested sequence. Response: cbind(n, y). Standard errors (Jaccard-adjusted for shared species) in parentheses. ^∗^*p <* 0.1; ^∗∗^*p <* 0.05; ^∗∗∗^*p <* 0.01.

**Table S4.**
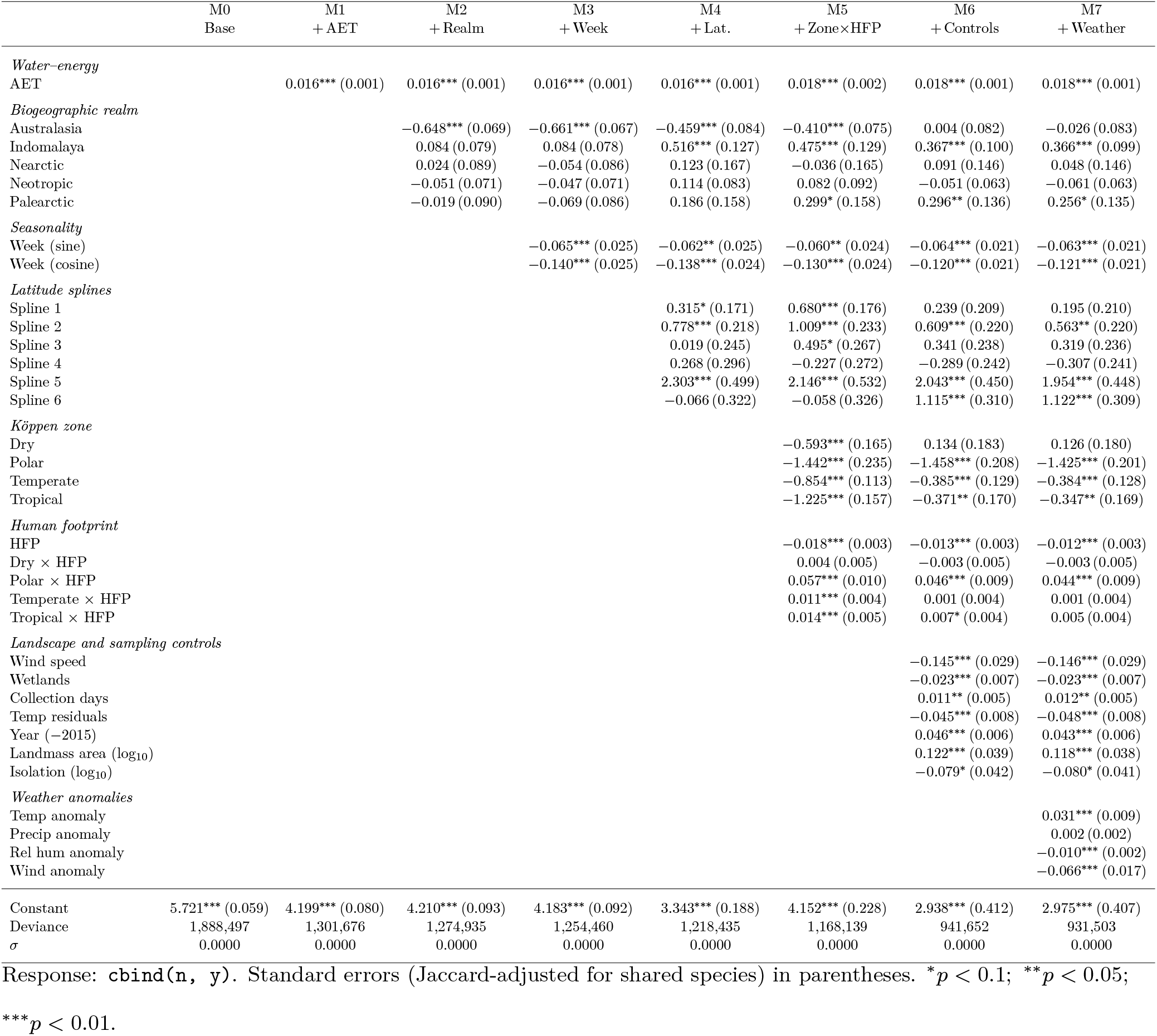
Stepwise Hubbell regression estimates (canonical link). Coefficients are added in nested blocks from the base intercept-only model (M0) to the full weather-anomaly model (M7). Realm and climate-zone coefficients are relative to the omitted reference category.

